# Metabolic Rewiring at the Pyruvate Node Drives Severe Pneumonia and T-Cell Suppression in Serotype 3 *Streptococcus pneumoniae* Infection

**DOI:** 10.64898/2026.02.13.705843

**Authors:** Kenichi Takeshita, Ana G. Jop Vidal, Jorge E. Vidal

## Abstract

**Background:** Serotype 3 (ST3) *Streptococcus pneumoniae* remains a major cause of invasive pneumococcal disease and pneumonia despite PCV13 introduction, in part due to potent immune evasion properties. The contribution of the pyruvate metabolic node (SpxB/LctO pathways) to ST3 pathogenesis is poorly defined.

**Methods:** We investigated oxygen-dependent fitness, virulence, and lung pathology in the ST3 strain WU2 and isogenic Δ*spxB*, Δl*ctO*, and Δ*spxB*Δ*lctO* mutants. In vitro growth was assessed under nasopharyngeal (21% O₂) and alveolar (14% O₂) conditions. Murine pneumonia models evaluated survival, bacterial burdens, histopathology (H&E, confocal microscopy), and lung transcriptomics (RNA-seq).

**Results:** ST3-strain exhibited a unique oxygen-sensitive growth defect at 21% O₂, alleviated by *spxB* deletion, indicating metabolic burden from pyruvate flux. In mice, wild-type WU2 caused high mortality with severe suppurative bronchopneumonia, alveolar consolidation, hemorrhage, and perivascular inflammation. The Δ*spxB* mutant accelerated lethality with enhanced distal lung damage, uncontrolled dissemination, and amplified inflammation. Wild-type infection uniquely induced targeted reorganization of bronchial epithelial membranes, forming prominent bacterium-laden blebs—host-derived membrane protrusions encapsulating intact pneumococci. These novel structures facilitated organized bacterial translocation into tissue without overt cytotoxicity and were largely absent in *spxB*-deficient mutants despite comparable lung burdens. RNA-seq analysis revealed SpxB-dependent suppression of T cell activation (e.g., *Rag1* and *Themis*) and acute inflammatory pathways, consistent with immune sequestration via blebs.

**Conclusions:** The SpxB-dependent pathway orchestrates ST3 virulence by enabling metabolic adaptation and driving bleb-mediated epithelial invasion and immune evasion in the lung. These bacterium-laden blebs represent a novel hallmark mechanism in pneumococcal pathogenesis, offering new insights and potential therapeutic targets.

## Introduction

*Streptococcus pneumoniae* (pneumococcus, Spn) remains a leading cause of community-acquired pneumonia and other lower respiratory infections, contributing to substantial global morbidity and mortality. According to the Global Burden of Disease Study 2023, Spn accounted for approximately 634,000 deaths (95% UI 565,000–721,000) attributable to lower respiratory infections, representing 25.3% of all such deaths [1]. Spn is classified into over 100 serotypes based on capsular polysaccharide antigenicity, with a subset responsible for most disease burden and targeted by pneumococcal conjugate vaccines (PCVs)[2]. Despite the inclusion of serotype 3 (ST3) in higher valent vaccines such as PCV13 (introduced in many national immunization programs since 2010), this serotype has persisted as a significant cause of invasive pneumococcal disease (IPD) in various settings [3–6]. ST3-associated IPD is frequently linked to severe clinical manifestations, including complicated pneumonia and empyema [7]. Notably, a high proportion of ST3 cases occur as vaccine breakthroughs in fully vaccinated children, including in certain post-pandemic surveillance cohorts [7–9].

ST3 strains characteristically form large, mucoid colonies when cultured on blood agar plates, a phenotype widely attributed to the production of an abundant capsular polysaccharide [10, 11]. The type 3 capsule is synthesized via a unique synthase-dependent pathway and consists of repeating →4)-β-D-GlcA-(1→4)-β-D-Glc-(1→ (cellobiuronic acid) units. This biosynthetic mechanism is associated with elevated capsule expression and partial extracellular release of polysaccharide, which contributes to the viscous, mucoid colony morphology[10, 11]. The virulence and relative persistence of ST3 despite inclusion in conjugate vaccines have been linked to several capsular properties, including increased capsule thickness and polysaccharide shedding, which may impair opsonophagocytic killing and facilitate immune evasion [12]. Additionally, ST3 strains exhibit a lower negative surface charge compared with other serotypes, potentially reducing complement deposition and further contributing to vaccine escape [13]. However, definitive causal evidence directly linking these phenotypic traits to enhanced virulence or reduced vaccine effectiveness remains incomplete, and additional mechanistic studies are required to fully elucidate their contributions.

ST3 pneumococci cause some of the most severe forms of pneumococcal lung disease. To the best of our knowledge, human lung histopathology from fatal ST3 infections is not publicly available. In contrast, murine models of pneumococcal pneumonia have provided detailed insights into the severe lung pathology elicited by ST3 strains. While pneumococcal infection in mice generally induces cytotoxicity and inflammatory infiltrates, ST3 infection elicits a distinct and markedly exacerbated pathological response. At 48 h post-inoculation, ST3 challenge is characterized by dense neutrophilic infiltrates, airway occlusion by suppurative exudates, extensive intrabronchial inflammation, perivascular edema and vascular leakage, necrotizing pleuritis, and steatitis [14, 15]. These features have been documented in C57BL/6 mice infected intranasally with ST3 strains such as NCTC 7978 or WU2, which develop expansive suppurative-to-necrotizing bronchopneumonia with peripheral lesion progression, pronounced peribronchial and perivascular inflammation, alveolar edema, and pleural involvement, reflecting disease severity that exceeds that observed in many other pneumococcal models [14, 15]. By comparison, murine pulmonary infection with non-ST3 pneumococcal strains, including TIGR4 (serotype 4), D39 (serotype 2), and EF3030 (serotype 19F), typically results in bronchiolar epithelial detachment, fibrotic alveolar changes, alveolar hemorrhage, and inflammatory cell infiltration[16–18]. However, extensive occlusion of bronchial airspaces by inflammatory exudates is not a prominent feature. In these models, pneumococci undergo decapsulation during bronchiolar invasion followed by re-encapsulation upon entry into the alveolar space, a process thought to mitigate toxicity from labile heme and other host-derived inflammatory mediators [16].

The vaccine escape properties of ST3-Spn have been extensively investigated, revealing mechanisms such as profuse shedding of its thick, loosely attached capsular polysaccharide, which likely neutralizes opsonizing antibodies, as well as emerging genomic variants with deletions in the capsular locus that further enhance immune evasion and persistence despite conjugate vaccine pressure. Pneumococcal lung infection elicits robust host transcriptional responses, including upregulation of oxidative stress-associated genes such as *Nos2* and *Hmox1*, reflecting cellular damage and redox imbalances observed in both *in vitro* and *in vivo* models [19–21]. This is accompanied by activation of the TLR–NF-κB innate immune pathway, driving proinflammatory cytokine production, neutrophil recruitment, and subsequent adaptive immunity [20, 22, 23]. Notably, host responses are serotype-dependent; microarray-based transcriptomic analyses in mouse pneumonia models have revealed that ST3 induces distinct profiles marked by strong expression of CXCR3-ligand chemokines (CXCL9, CXCL10, and CXCL11) compared to other serotypes [24]. Furthermore, IL-17-mediated neutrophil uptake and killing of ST3 were impaired compared to serotype 4 strain [25]. However, dual RNA-seq studies in C57BL/6 mice intranasally infected with the ST3 strain SRL1 have shown downregulation of neutrophil antimicrobial functions, including reduced expression of neutrophil elastase and other granule proteins. This potentially impairs bacterial clearance despite massive influx of neutrophils, contributing to persistent infection and heightened pathology[26].

Glycolysis, funneling into the pyruvate node, represents a primary metabolic pathway supporting Spn during lung pathogenesis. Unlike many other streptococcal species, Spn predominantly converts pyruvate to lactate via lactate dehydrogenase (Ldh) to regenerate NAD⁺. Subsequent re-oxidation of lactate to pyruvate is mediated by lactate oxidase (LctO), an O₂-dependent reaction that irreversibly produces hydrogen peroxide (H₂O₂) as a byproduct. Concurrently, pyruvate is oxidized by pyruvate oxidase (SpxB), a flavin mononucleotide-dependent enzyme that also uses O₂ as the electron acceptor, yielding acetyl phosphate, ATP (via acetate kinase, AckA), and H₂O₂, which is considered the major source of endogenous oxidant in pneumococcal cultures. Both SpxB and LctO reactions are strictly oxygen-dependent. Molecular O₂ is readily accessible in the aerated alveolar spaces of the lungs, even amid infection-induced inflammation, thereby enabling sustained activity of this pathway. The essential nature of acetate kinase and therefore the importance of this pathway during the intracellular stage are underscored by the inability to stably delete the *ackA* gene without compensatory mutations.

During the intracellular phase of infection, critical for invasive pneumococcal disease, this glycolytic-pyruvate node pathway likely supplies Spn with essential nutrients while facilitating immune evasion, particularly in ST3 strains. Emerging evidence indicates that Spn manipulates host cell metabolism, including glycolysis and oxidative phosphorylation, to access intracellular pyruvate pools; however, the precise molecular mechanisms remain incompletely understood. In this study, we investigated the contribution of pyruvate node enzymes to the pathogenesis of ST3 strains. Our findings reveal that the concerted activity of these enzymes exerts substantial control over the host immune response, as reflected by heightened pulmonary pathogenesis, disrupted invasion dynamics, and enhanced stimulation of specific T-cell responses in the absence of key pyruvate node components.

## Methods

### Bacterial strains media, and regents

All Spn strains utilized in this study are listed in the Table 1.

**Table 1.** Strains utilized in this study.

| Strain | Characteristics | Reference |
| --- | --- | --- |
| WU2 | Reference strain, whole genome sequenced, capsular serotype 3 | This study |
| WU2Δ*spxB* | WU2 with an insertionally inactivated *spxB::ermB* (Ery<sup>r</sup>) | This study |
| WU2Δ*lctO* | WU2 with an insertionally inactivated *lctO::aad9* (Spc<sup>r</sup>) | This study |
| WU2Δ*spxB*Δ*lctO* | WU2 with a deleted *spxB* and *lctO* gene by insertion-deletion with an erythromycin and spectinomycin cassette, respectively | This study |
| WU2Δ*spxB*Ω*spxB* | complemented with the gene *spxB* and its promoter | This study |
| TIGR4 | Reference strain, whole genome sequenced, capsular serotype 4 | [27] |
| TIGR4Δ*spxB* | TIGR4 with an insertionally inactivated *spxB::ermB* (Ery<sup>r</sup>) | [28] |
| TIGR4Δ*lctO* | TIGR4 with an insertionally inactivated *lctO::aad9* (Spc<sup>r</sup>) | [29] |
| TIGR4Δ*spxB*Δ*lctO* | TIGR4 with a deleted *spxB* and *lctO* gene by insertion-deletion with an erythromycin and spectinomycin cassette, respectively | [29] |
| D39 | Avery strain, clinical isolate, capsular serotype 2 | [30] |
| EF3030 | Clinical isolate, capsular serotype 19F | [31] |

Strains were inoculated on cultured blood agar plate (BAP) containing 10 μg/ml gentamicin (Sigma-Aldrich) or the appropriate antibiotics (for the isogenic mutants) from frozen stocks stored in skim milk, tryptone, glucose, and glycerin. These cultures were incubated at 37°C for 24 hours in 21% O_2_ and 5% CO_2_ atmosphere, followed by replating on BAP without antibiotics (Fisher Scientific) and incubating overnight under the same conditions) Other reagents used and sources were the following: erythromycin (Sigma-Aldrich), spectinomycin (Sigma-Aldrich), chloramphenicol (Sigma-Aldrich). Todd-Hewitt broth (Becton Dickinson), yeast extract (Becton Dickinson).

### Preparation of inoculum for *in vitro* and for animal experiments

Inoculum for the *in vitro* studies was prepared essentially as previously described with minor modifications. Briefly, the overnight BAP culture of the Spn strain was used to prepare a bacterial suspension in sterile Todd-Hewitt broth supplemented with 0.5% yeast extract (THY). This fresh bacterial suspension was inoculated to a final optical density at 600 nm (OD_600_) of ∼0.1 corresponding to ∼5.15 × 10⁸ CFU/mL. Unless otherwise noted, all *in vitro* experiments described below were inoculated with fresh inocula, i.e., prepared right before the start of the experiment.

To prepare inoculant for mice studies, the Spn strain was inoculated in THY broth supplemented with 20% fetal bovine serum (FBS; Corning); this culture was grown for 4 h (i.e., early log phase) at 37°C in a 5% CO_2_ atmosphere; the growth was monitored using a spectrophotometer Bio-Rad SmartSpec Plus. After this incubation time, the culture was centrifuged at 10,000 rpm for 10 min and the pellet was resuspended in 10 ml phosphate buffered saline (PBS, Fisher Scientific) (pH=7.4) and washed once. The washed bacterial pellet was resuspended in PBS containing 10% (v/v) glycerol and stored at -80ᵒC until it was used. Aliquots of inocula were thawed from each batch to obtain the density by dilution and plating. All inocula were used within two months.

### Preparation of WU2 isogenic mutant derivatives

Isogenic mutants in the ST3 strain WU2 were constructed by natural transformation using donor DNA from previously generated TIGR4 (serotype 4) mutants, leveraging the high sequence homology at the *spxB* and *lctO* loci. Genomic DNA from donor strain TIGR4Δ*spxB* (with *spxB* insertionally deleted and replaced by an erythromycin resistance cassette [erm]) and TIGR4Δ*lctO* (with *lctO* insertionally inactivated by a spectinomycin resistance cassette [spc]) was extracted using the QIAamp DNA Mini Kit (Qiagen). For the double mutant, DNA from TIGR4Δ*spxB*Δ*lctO* (carrying both cassettes) was used. Competent WU2 cells were prepared and transformed using a standard natural competence protocol for Spn [32]. Briefly, cells were grown to early exponential phase in THY, rendered competent by transfer to competence medium, and incubated with ∼1 µg donor DNA for 2 h at 37°C to allow uptake and homologous recombination. Transformants were selected on BAP supplemented with erythromycin (1 µg/mL) for Δ*spxB*, spectinomycin (100 µg/mL) for Δ*lctO*, or both antibiotics for the double mutant Δ*spxB*Δ*lctO*. Insertional inactivation was confirmed by PCR using primers spxB-confirmation1 and spxB-confirmation2 for *spxB*, and analogous primers for *lctO* (Table 2).

**Table 2.**
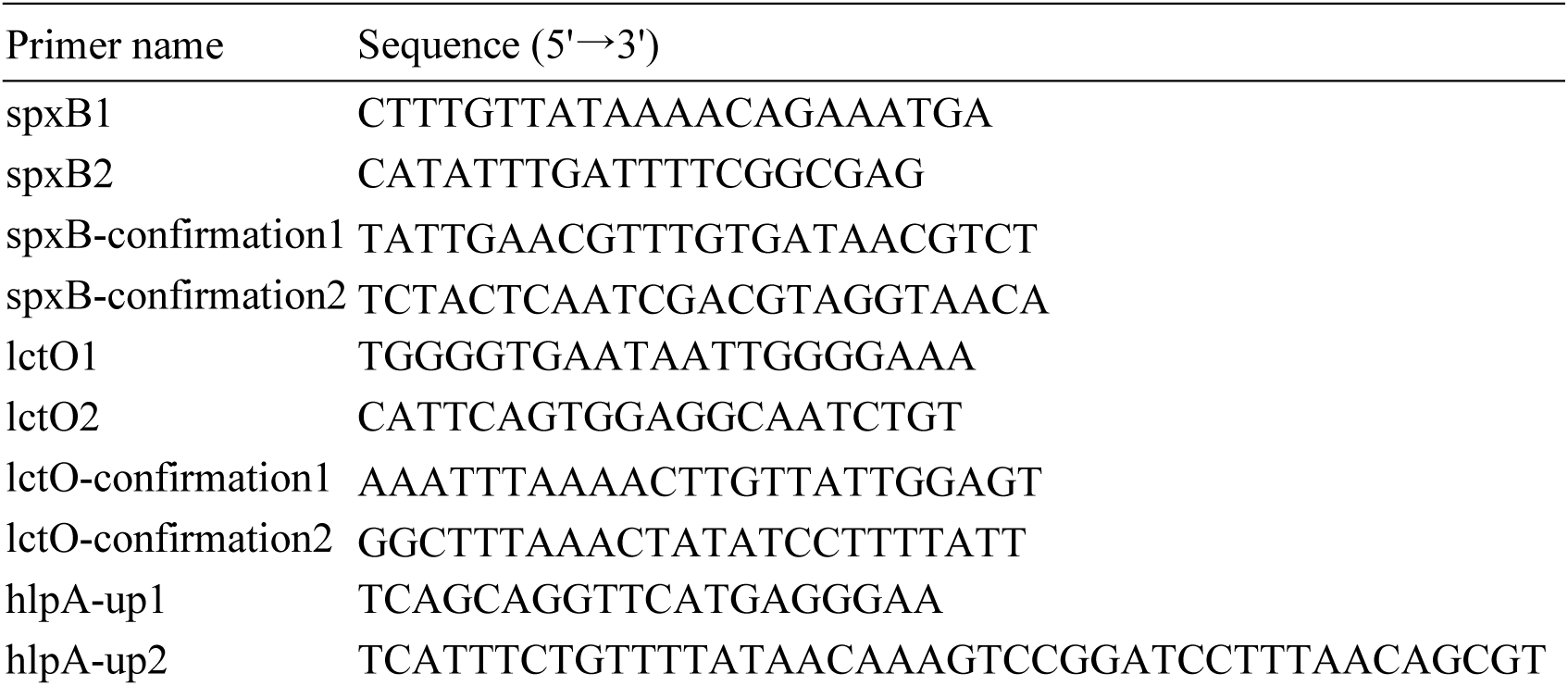
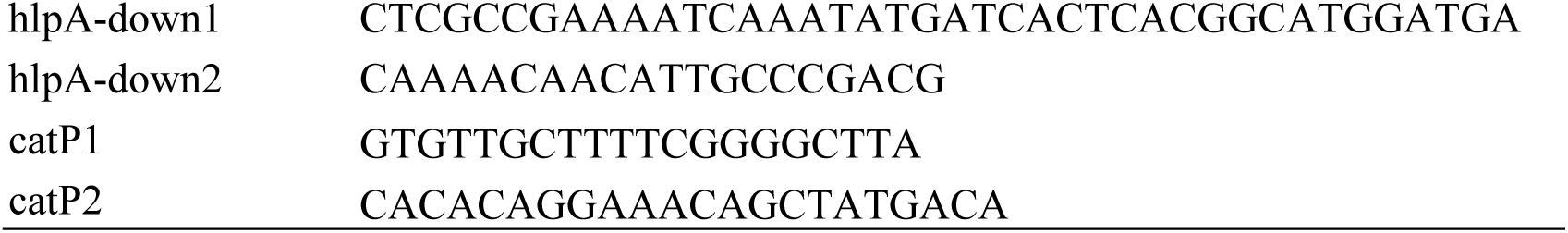
Oligonucleotides designed and used in this study.

Mutant candidates were further screened by growth kinetic analysis. Representative clones were selected for whole-genome sequencing (WGS) to verify the intended mutation(s), confirm clean allelic replacement, and rule out unintended secondary insertions, deletions, or off-target mutations. WGS was performed at the SeqCenter (Before known as MiGS Microbial Genome Sequencing Center) using the NextSeq 2000 platform.

### Complementation of *spxB* in WU2Δ*spxB*

To complement the gene *spxB* in WU2Δ*spxB*, a “neutral” chromosomal locus within the *hlpA* gene (SP1113) was selected, as insertions at this site do not affect virulence [33]. The *spxB*-complemented fragment was generated by amplifying an upstream region containing part of the *hlpA* gene using DNA purified from WU2 as a template and primers, hlpA-up1 and hlpA-up2 (Table 2). A PCR fragment containing the *spxB* gene and its promoter was amplified from WU2 genomic DNA using primers, spxB1 and spxB2. A downstream fragment comprising the *catP* gene, encoding chloramphenicol resistance, and a fragment of open reading frame SP1114 was amplified from strain JWV500 DNA using primers, hlpA-down1 and hlpA-down2. All PCR products were purified using a GeneJET PCR Purification Kit (Thermo scientific), and subsequently ligated by splicing overlap extension PCR with primers hlpA-up1 and hlpA-down2. The resulting PCR-ligated product was purified as described above and used to transform WU2Δ*spxB* following standard procedure [32]. Transformants were selected on blood agar plates containing chloramphenicol (3.5 µg/mL). Candidate clones were screened by PCR using primers flanking the insertion site to confirm successful integration (Table 2). Positive clones were further validated by growth kinetics under controlled conditions and by WGS to verify the integrity of the complementation construct and the absence of unintended mutations.

### Growth kinetics

THY was inoculated with Spn strains and their corresponding mutant derivatives in a 24-well plate (Genesee Scientific). Plates were incubated in a microplate reader (Agilent BioTek Synergy H1) at 37°C under some specific O_2_ and CO_2_ conditions of atmosphere. The OD_600_ was measured every 20 min for 24 h to generate growth curves. Uninfected THY was always incubated with the samples as the blank. For experiments in the presence of exogenous catalase, 100 units (U) of filter-sterilized liquid catalase (Sigma-Aldrich) was added in 1mL of bacteria-contained THY before the incubation.

### Cell cultures of human respiratory cells

Human bronchial Calu-3 cells (ATCC, HTB-55) were used for the *in-vitro* cell culture adhesion models. Calu-3 cells were cultured in Eagle’s Minimum Essential Medium (EMEM, ATCC) supplemented with 10% FBS and 100 U/mL of penicillin-streptomycin (Gibco) in 75 cm^2^ flasks. Cells were incubated at 37°C with 5% CO_2_ atmosphere and supplemented with fresh medium three times weekly and passaged to a new flask once weekly or when cells reached ∼100% confluency by trypsinization with Trypsin-EDTA (0.25%, ATCC), phenol red (Gibco) and seeded in the experiment specific device.

### Quantification of hydrogen peroxide production

H₂O₂ production by the strains was quantified using the Amplex Red Hydrogen Peroxide Assay kit (Invitrogen) in a 96-well plate format according to the manufacturer’s instructions. Briefly, strains were incubated in 6 well plates containing 2 mL of fresh THY media at 37 °C in a 5 % CO_2_ atmosphere for 4 h. Following incubation, culture supernatants were collected and centrifuged at 4 °C for 5 min at 15,000 rpm. The clarified supernatants were transferred to fresh tubes and kept on ice for no longer that 30 min prior to analysis. Samples were diluted in 1x reaction buffer provided with the Amplex Red H_2_O_2_ assay kit, and H_2_O_2_ levels were measured using a microplate reader (Agilent BioTek Synergy H1).

### Qualitative Detection of Hydrogen Peroxide production

Qualitative assessment of H₂O₂ production was performed using Prussian blue agar plate as previously described [34]. Prior to inoculation, 200 µL of commercially available human serum (MP Biomedicals) was added to the plates to supplement bacterial growth. Strains were then inoculated onto the prepared plates and incubated under the same conditions. H₂O₂ production was indicated by the formation of a blue coloration on the agar surface.

### Ethics Statement

All experiments involving animals were performed with prior approval of and in accordance with protocol 2022-1245 which was reviewed and approved by the University of Mississippi Medical Center (UMMC) Animal Care Committee. UMMC laboratory animal facilities have been fully accredited by the American Association for Accreditation of Laboratory Animal Care. Procedures were performed according to the institutional policies, Animal Welfare Act, National Institutes of Health (NIH) guidelines, and American Veterinary Medical Association guidelines on euthanasia.

### Animal study information

Four-week-old male C57BL/6 mice were obtained from Charles River Laboratory (Wilmington, MA, USA) allowed to acclimate for one week prior to the experimental challenge. Mice were maintained under standard housing conditions and provided food and water ad libitum throughout the study. The mice were infected intranasally with 1×10^8^ CFU of Spn resuspended in 30 μL of PBS. The mice inoculated with the same strain housed in the same cage. The behavior and the weight of the mice were monitored twice a day. Mice were humanly euthanized via inhalation of 5 % isoflurane at the indicated period, or when they lose over 15 % of their body weight, or when mice were non-responsive to manual stimulation, and/or if they show signs of illness such as ruffled fur, intermittent hunching, and exhibiting respiratory distress.

Blood, nasopharynx, trachea, and lung tissues were collected in post-euthanasia. Blood was collected in a tube and immediately added with THY broth added with 10 % glycerol, and used for bacterial counting making dilution and plating onto BAP containing gentamicin. Nasopharyngeal tissue and tracheal tissue collected in THY broth added with 10 % glycerol and stored in -80 ℃. These samples were homogenized the other days and plated onto BAP containing gentamicin to count bacterial density. Lung tissue was split half and one was analyzed bacterial density as the nasopharynx and trachea, and the other were collected in 4 % paraformaldehyde (PFA) and immediately stored at -80 °C for histological analysis and confocal microscopy.

### IL-6 quantification in the mouse lung tissue by enzyme linked immunosorbent assay (ELISA)

The 80 uL of homogenized lung samples were centrifuged at 4000 g for 10 min to collect supernatants. The samples were diluted with the kit-contained diluent if needed. IL-6 concentrations of the lung tissue were measured in a 96 well plate using Mouse IL-6 Uncoated ELISA kit (Invitrogen) according to the manufacturer’s instructions.

### Histopathology

PFA-fixed lung tissues were paraffin-embedded, sectioned (∼5 μm) and stained with hematoxylin and eosin (H&E) at the UMMC histology core facility. Brightfield images were acquired using Olympus microscope.

For immunofluorescence analysis, paraffin-embedded lung sections were deparaffinized and washed with PBS, followed by blocking with 2 % bovine serum albumin (BSA, Roche Diagnostics) for 30 min at room temperature. Sections were then incubated for 2 h at room temperature with wheat germ agglutinin (WGA) conjugated to Alexa Fluor 488 (Molecular Probes) (5 µg/mL), and an Alexa Fluor 555-conjugated anti-ST3 capsular polysaccharide antibody (Serum Institute, Denmark) (40 μg/mL). After incubation, sections were washed with PBS, air-dried and mounted using ProLong Diamond Antifade mountant containing DAPI (Molecular Probes).

### Quantification of high-resolution confocal micrographs

Confocal z-stack micrographs were analyzed using Imaris software (version 10.1.0, 64-bit; Oxford Instruments). Regions of interest (ROIs) were manually defined within the z-stacks to quantify the surface area (µm²) of host plasma membranes (WGA-AF488 channel), cell nuclei (DAPI channel), and pneumococcal capsule (Alexa Fluor 555 anti-serotype 3 channel). For 3D visualization and spatial analysis, surface masks were generated for each channel using Imaris surface reconstruction algorithms. The shortest distance from membrane surfaces to nuclear surfaces was calculated using built-in distance transformation tools. Distances were binary classified into two categories: ≤5 µm (proximal/interacting) and >5 µm (distal). Areas were normalized as the percentage of total membrane or capsule area falling within ≤5 µm of nuclei and presented as a proxy for epithelial cytotoxicity or membrane disruption.

To assess co-localization between capsule and membrane signals, overlap volume analysis was performed using the Imaris Coloc module with a threshold of 0.1 µm; voxels within this threshold were considered true co-localization. For simplicity and standardization across experiments, all quantitative outputs (area percentages, distance classifications, and co-localization volumes) were converted to percentages of the total relevant signal within the ROI. Data are presented as mean ± SE unless otherwise specified, with statistical comparisons performed as indicated in the corresponding figure legends.

### RNA-sequence of the mouse tissue

C57BL/6 4-week-old male mice were infected as described above. The lung samples were collected post-euthanasia 48 h post-infection, flash frozen and stored at -80°C. The following procedures were performed by the UMMC Molecular and Genomics Core Facility (MGCF) (www.umc.edu/genomicscore). The tissue was homogenized in 1 mL TRIzol per 50-100 mg tissue with a homogenizing machine and centrifuged to collect supernatant in fresh RNase-free tube. RNA was extracted using the Pure Link RNA Mini Kit from archived samples (Invitrogen) according to the manufacturer’s instructions and assessed for quality control parameters of minimum concentration and fidelity (i.e., 18S and 28S bands, RIS > 8) through the MGCF. Libraries were prepared for RNA sequencing per detailed protocols from the Illumina mRNA stranded kit. Briefly, a pooled library was prepared from using n = 12-24 samples per group using TruSeq mRNA Stranded Library Prep Kit (Set-A-indexes), quantified with the Qubit Fluorometer (Invitrogen), and assessed for quality and size using Qiagen QIAxcel Advanced System or Agilent TapeStation. The library was sequenced using the NextSeq 2000 using P3 (200 cycles, paired end 100 bp) on the Illumina NextSeq 2000 platform. Sequenced reads were assessed for quality using the Illumina BaseSpace Cloud Computing Platform and/or custom analysis pipeline, and FASTQ sequence files were aligned reads to the reference genome (e.g., Ensembl/Hg38, Rnor_6.0) using DRAGEN RNA Pipeline Application (v.3.7.5) or custom analysis pipe-lines developed through MGCF. Differential expression was determined using DRAGEN Differential Expression Application (v.3.10.4) of DESeq2. differentially expressed genes (DEGs) were identified using cutoffs of log2 (fold change) ≥ 1 (2-fold), with an adjusted P-value < 0.05. Data were deposited into NCBI GEO data under accession number (It will be provided during revisions).

To elucidate the biological implications of Spn infection, functional enrichment analysis was performed on DEGs using DAVID (v2023q1) and g:Profiler (v.e109_eg56_p17). DAVID and g:Profiler were used to identify enriched GO terms across three categories: Biological Process (BP), Molecular Function (MF), and Cellular Component (CC). GO-BP analysis revealed significant enrichment of terms related to “oxidoreductase activity, acting on paired donors, with incorporation or reduction of molecular oxygen” (GO:0016705, padj = 2.845×10^−3^), consistent with Spn-H₂O₂-induced oxidation of the host tissues. GO-BP analysis also highlighted “T cell activation” (GO:0042110, padj = 4.990×10^−4^), which can lead to immune evasion. To focus on differences in gene expression between wild-type WU2 infected tissue and WU2Δ*spxB*-infected tissue, the genes that met the criteria of padj < 0.05 and log2 fold change ≥ 1 in this comparative dataset were extracted and compared the differential gene expression of these genes in the dataset relative to mock-infected tissue.

## Statistical Analysis

We performed one-way analysis of variance (ANOVA) followed by Šidák’s multiple comparison test or Dunnett’s multiple-comparisons test when two or more groups were involved. All analysis was performed using GraphPad Prism software (version 10.6.1).

## Results

### Oxygen-dependent growth dynamics in serotype 3 *Streptococcus pneumoniae*

During colonization and dissemination in the respiratory tract, Spn is exposed to gradients in oxygen availability. The nasopharynx typically presents near-atmospheric oxygen levels (∼21% O₂), whereas the lower airways and alveoli exhibit reduced oxygen tension (∼14% O₂), often microaerophilic. To assess the influence of oxygen on Spn fitness, we compared the growth kinetics of the reference ST3 strain, WU2 with representative vaccine and non-vaccine serotypes—D39 (serotype 2), TIGR4 (serotype 4), and EF3030 (serotype 19F)—under conditions mimicking nasopharyngeal (21% O₂) and lower respiratory tract/lung (14% O₂) environments over 24 h.

At 21% O₂, WU2 displayed markedly impaired growth relative to the other serotypes (Fig. 1A). This phenotype was evident as a prolonged lag phase, decreased maximum growth rate (V_max), lower peak OD₆₀₀, and extended time to achieve V_max (T at V_max) compared with D39, TIGR4, and EF3030 (Fig. S1A). In contrast, at 14% O₂, WU2 exhibited enhanced growth kinetics, surpassing its performance at 21% O₂ and outperforming the comparator serotypes (Fig. 1B, 1C; Fig. S1B). Specifically, WU2 showed superior growth parameters relative to TIGR4 and EF3030 under these reduced-oxygen conditions (Fig. 1B; Fig. S1C). This oxygen-dependent growth defect was unique to WU2 and clinical ST3-Spn isolates (not shown) among the serotypes examined, indicating a specific sensitivity of ST3 to higher oxygen concentrations typical of the nasopharyngeal niche.

**Figure 1.**
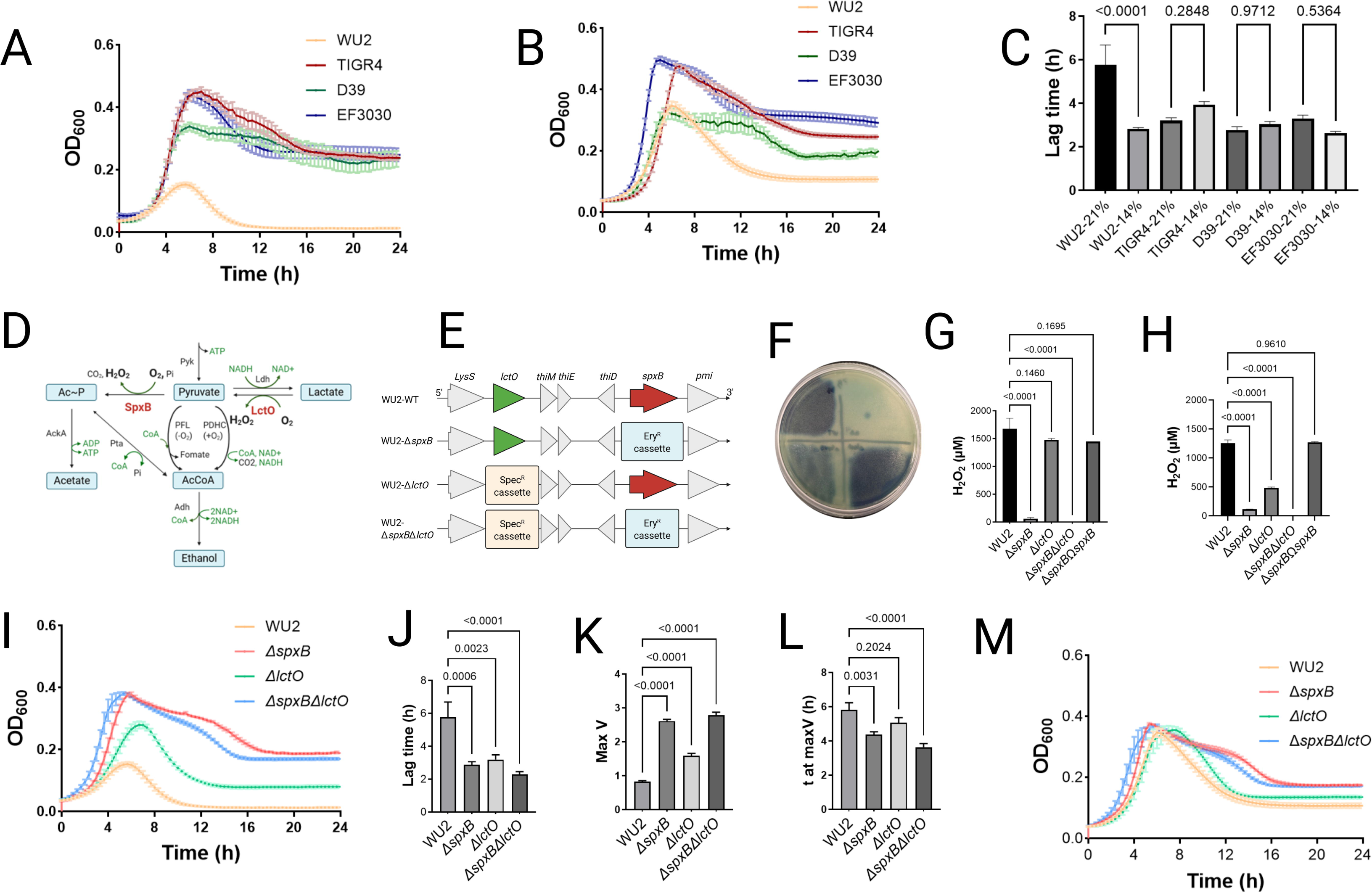
Oxygen-dependent growth dynamics in serotype 3 *Streptococcus pneumoniae* and the impact of pyruvate node metabolism on the fitness. (A, B) The growth kinetics of WU2, TIGR4, D39, and EF3030 in the nasopharyngeal oxygen environment (A: 21% O_2_ with 5% CO_2_) or in the lung oxygen environment (B: 14% O_2_ with 5% CO_2_). The strains were cultured in Todd-Hewitt broth supplemented with 0.5% yeast extract (THY) at 37°C, and OD₆₀₀ was measured every 20 min and plotted. (C) The comparison of lag time (h) of the growth curves between the different oxygen environments. (D) Pyruvate node in the pneumococcal metabolic pathway. (E) Genomic context of the *spxB* and *lctO* genes in WU2 and insertional deletions within *spxB* (red arrow)/*<u>lctO</u>* (green arrow) with an erythromycin (Ery^R^)/ spectinomycin (Spec^R^) resistance cassette. (F-H) Hydrogen peroxide production of wild-type WU2 and the isogenic mutants. The strains were (F) incubated for 24 h onto Prussian Blue agar plate (WU2; left upper, WU2Δ*spxB*; left lower, WU2Δ*lctO*; right lower, WU2Δ*spxB*Δ*lctO*; right upper) under an ambient oxygen environment or (G, H) inoculated in THY and incubated at 37°C for 4 h under (G) the nasopharyngeal oxygen environment (21% O_2_ with 5% CO_2_) or (H) the lung oxygen environment (14% O_2_ with 5% CO_2_), and hydrogen peroxide was quantified with AmplexRed^®^. (I) The growth kinetics of WU2, WU2Δ*spxB*, WU2Δ*lctO*, and WU2Δ*spxB*Δ*lctO* in the nasopharyngeal oxygen environment (21% O_2_ with 5% CO_2_). (J-L) The comparison of growth parameters of the WU2 and the mutants. Bars represent mean values of (J) lag time (h), (K) maximum growth rate (Max V, 1/h), and (L) time to maximum growth rate (t at Max V, h). (M) The growth kinetics of WU2, WU2Δ*spxB*, WU2Δ*lctO*, and WU2Δ*spxB*Δ*lctO* in the lung oxygen environment (14% O_2_ with 5% CO_2_). Data in all panels are shown as mean ± SE. Statistical analyses in panels (C, G, H, J, K, and L) were performed using one-way ANOVA with Šidák’s multiple comparisons test.

### Impact of pyruvate node metabolism on serotype 3 *Streptococcus pneumoniae* fitness across oxygen gradients and under oxidative stress

Serotype 3 strain WU2 exhibits pronounced sensitivity to oxidative stress, and oxygen availability modulates flux through the pyruvate node via the enzymatic activities of SpxB and LctO, which in turn influence key pneumococcal metabolic pathways (Fig. 1D). To investigate the role of these enzymes, we constructed isogenic mutants by targeted deletion of *spxB* and *lctO* in the WU2 background. Successful gene inactivation was confirmed by whole-genome sequencing (Fig. 1E).

We next evaluated the fitness consequences of disrupted pyruvate node metabolism by measuring growth kinetics and extracellular H₂O₂ production in wild-type WU2 and its mutant derivatives (Δ*spxB*, Δ*lctO*, Δ*spxB*Δ*lctO*, and the *spxB*-complemented strain) under nasopharyngeal-mimetic (21 % O₂) and lower respiratory tract-mimetic (14 % O₂) conditions. Under nasopharyngeal conditions, extracellular H₂O₂ production was markedly reduced in the Δ*spxB* and Δs*pxB*Δ*lctO* mutants compared to wild-type WU2, the Δ*lctO* single mutant, and the complemented strain, as quantified by Prussian blue assay and Amplex Red absorbance (Fig. 1F, G).

Under lung-mimetic conditions (14% O₂), H₂O₂ production by wild-type WU2 was already reduced by ∼15 % relative to 21 % O₂, whereas the Δ*lctO* mutant exhibited a substantially greater reduction (∼71 %; Fig. S2). The substantial reduction in H₂O₂ production observed in the Δ*lctO* mutant under lung-mimetic conditions suggests tightly controlled regulation of pyruvate node metabolism in the lower airways, likely optimizing fitness in this physiologically relevant oxygen gradient. At 21% O₂, mutations in *spxB* alone or in combination with *lctO* conferred the greatest fitness advantage, manifested as a shortened lag phase, elevated maximum growth rate (Max V), and reduced time to Max V (t at Max V) (Fig. 1I–L). Complementation of *spxB* by ectopic insertion into a neutral chromosomal locus restored growth parameters to levels comparable to wild-type WU2 (Fig. S3A–E). These aerobic growth defects in the wild-type WU2 were partially or fully rescued by supplementation with catalase (100 U/mL), confirming that excess endogenous H₂O₂ accumulation impairs growth under high-oxygen conditions (Fig. S3J).

Under 14 % O₂, growth kinetics of wild-type WU2 and all mutant derivatives were largely equivalent; however, wild-type WU2 exhibited a significantly lower Max V compared with the mutants (Fig. 1M; Fig. S3F–I). These results indicate that SpxB-dependent pyruvate flux impairs WU2 fitness in high-oxygen environments. In contrast, this pathway is disadvantageous for the host under lower-oxygen conditions typical of the lower respiratory tract, where ST3 pneumococci exhibit optimal growth, consistent with the bacterium’s adaptation to microaerophilic/hypoxic niches during pulmonary invasion.

### Virulence and attenuation of serotype 3 *S. pneumoniae* in a murine pneumonia model

To evaluate the contributions of pyruvate metabolism to fitness and virulence of ST3-Spn during respiratory tract infection, five week-old male C57BL/6 mice were intranasally inoculated with wild-type strain WU2 or its isogenic mutants: WU2Δ*spxB*, WU2Δ*lctO*, or WU2Δ*spxB*Δ*lctO*. PBS-inoculated mice served as mock-infected controls. Outcomes monitored over 96 h post-infection included survival, bacterial burdens in the nasopharynx/nasal septum (upper respiratory tract), trachea, lungs, and blood, and lung histopathology.

Infection with wild-type WU2 resulted in 75 % mortality by 96 h, with a median survival time of 4 days (Fig. 2A). The Δ*spxB* mutant induced equivalent overall lethality (75 % mortality) but significantly accelerated disease progression, shortening median survival to 2.5 days (Fig. 2A). In contrast, the Δ*lctO* mutant exhibited moderate attenuation (67 % mortality; median survival 4 days), whereas the double mutant (Δs*pxB*Δ*lctO*) partially suppressed the accelerated lethality of Δ*spxB*, yielding 50 % mortality and a median survival time of 4 days (Fig. 2A). These results indicate that disruption of SpxB-dependent pyruvate metabolism impairs host tolerance and accelerates disease progression, whereas concurrent inactivation of LctO mitigates this effect, restoring survival kinetics comparable to wild-type.

**Fig. 2.**
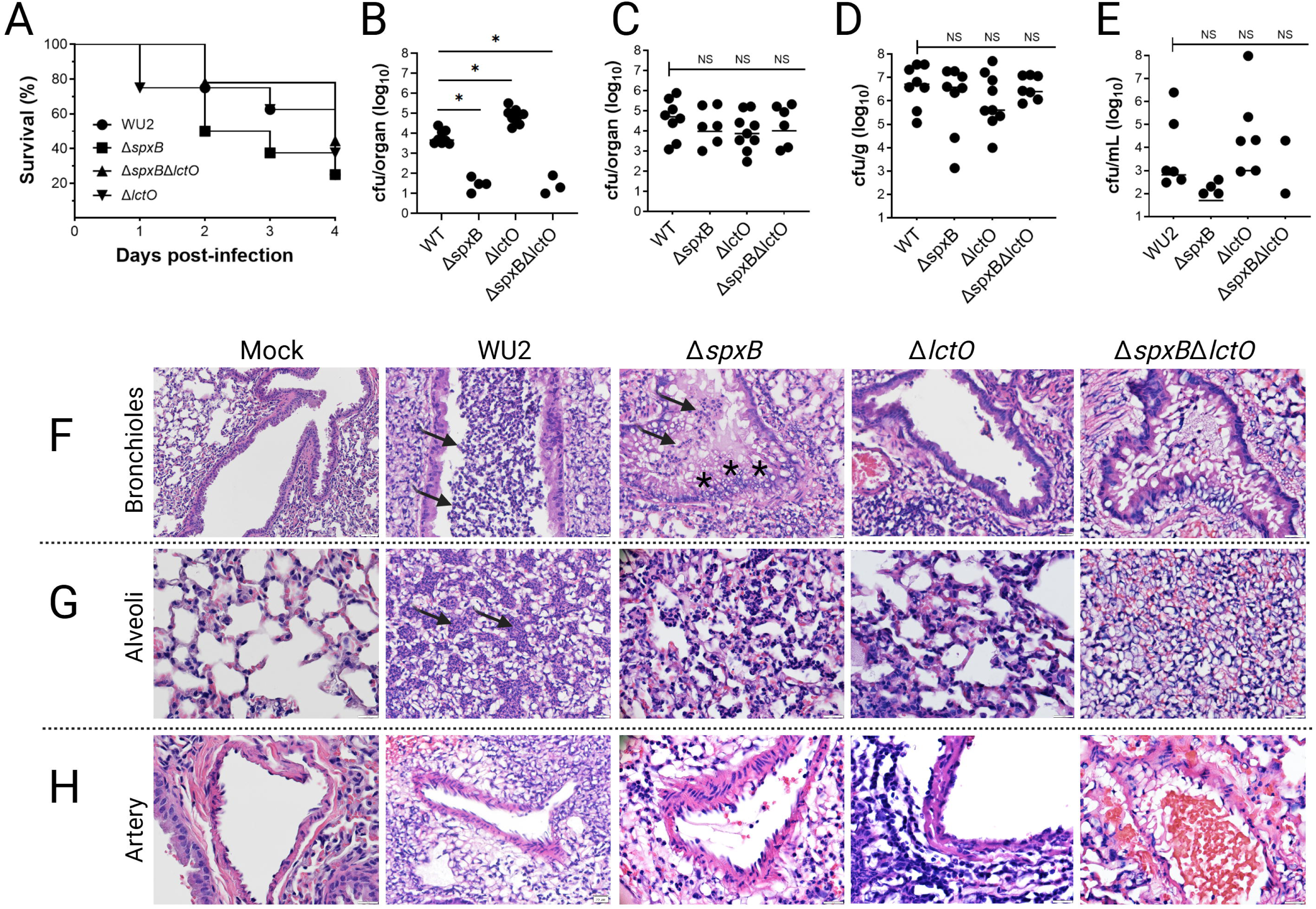
Virulence and attenuation of serotype 3 *S. pneumoniae* in a murine pneumonia model. (A) Kaplan-Meier survival curves of C57BL/6 male mice (n=8 per group) following intranasal infection with wild-type WU2 (WT), Δs*pxB*, Δ*lctO*, or Δ*spxB*Δ*lctO* (1×10^8^ CFU inoculum). Mice were monitored for 4 days post-infection. (B-E) The tissues and blood were harvested and homogenized at endpoint to quantify the bacterial burden. Bacterial burden of the (B) nasopharynx/nasal septum, (C) Trachea, (D) lung, and (E) Blood. Statistical comparisons of bacterial density were performed using Kruskal-Wallis test with Dunn’s multiple-comparisons test. ns, not significant. \**p*<0.05. (F–H) Representative hematoxylin and eosin-stained lung sections from mock-infected (PBS) or infected mice at the experimental endpoint. (F) Bronchioles: WT and Δ*spxB* show similar epithelial swelling (asterisk), luminal narrowing, and dense inflammatory cell infiltration (arrow; primarily neutrophils and macrophages); milder changes are observed in Δ*lctO* and especially Δ*spxB*Δ*lctO*. (G) Alveoli: WT and Δ*spxB* exhibit comparable alveolar consolidation, septal thickening, edema, and heavy inflammatory infiltrates (arrow); pathology is substantially reduced in the double mutant, with more preserved alveolar spaces in Δ*lctO*. (H) Pulmonary arteries/arterioles: WT and Δ*spxB* display similar perivascular inflammation, cuffing, and cellular exudates; attenuation is most evident in Δ*spxB*Δ*lctO*, with milder vascular involvement in Δ*lctO*. Magnification is shown in each panel. Images are representative of n=3–5 mice per group.

Bacterial burdens were quantified in the respiratory tract and blood at euthanasia or at 96 h in surviving animals. In the upper respiratory tract (nasopharynx/nasal septum), both wild-type WU2 and WU2Δ*lctO* achieved robust colonization in all animals, with median recoveries of 8.6 × 10³ CFU and 1.0 × 10⁵ CFU per organ, respectively, at euthanasia or at 96 h in survivors (Fig. 2B). By comparison, colonization by WU2Δ*spxB* and WU2Δ*spxB*Δ*lctO* was severely compromised: viable bacteria were recovered from only 4/8 and 3/8 mice, respectively, with median burdens in colonized animals below 2 × 10¹ CFU/organ—a reduction of >2–4 logs relative to WU2 and WU2Δ*lctO* (P = 0.0078, Mann–Whitney U test; Fig. 2B). Tracheal colonization showed a similar pattern: WU2 and WU2Δ*lctO* colonized all mice, whereas WU2Δ*spxB* and WU2Δ*spxB*Δ*lctO* colonized only 6 of 8 mice (75%) per group (Fig. 2C). Among colonized mice, however, tracheal bacterial densities were comparable across strains (median 10⁴–10⁵ CFU/organ), suggesting that the *spxB* defect primarily impairs initial establishment rather than subsequent replication at this site (Fig. 2C).

In the lungs, wild-type WU2 reached a median burden of 1.34 × 10⁷ CFU/g of tissue, whereas all mutant strains showed lower median burdens (<8.69 × 10⁶ CFU/g); these differences were not statistically significant (Fig. 2D). Systemic dissemination was more markedly impaired: bacteremia occurred in 75% of mice infected with WU2 or WU2Δ*lctO*, compared with 50% for WU2Δ*spxB* and 25% for WU2Δ*spxB*Δ*lctO*. Among bacteremic animals, blood CFU counts were similar across groups (P > 0.068, Mann–Whitney U test; Fig. 2E).

Collectively, these findings underscore the essential role of SpxB-dependent pyruvate metabolism in establishing upper airway colonization and facilitating progression to pneumonia and bacteremia in ST3-Spn, with LctO providing a compensatory mechanism within the pyruvate metabolic node in the absence of SpxB.

### Differential Lung Pathology Across Wild-Type WU2 and Pyruvate Node Mutants

Histological examination of lung tissues from different anatomical regions (bronchioles, alveoli, and arterioles/vessels) revealed marked differences in pathology (Fig. 2F-H). In mock-infected control mice, lung architecture was normal across all regions: bronchioles showed intact pseudostratified columnar epithelium with open lumens and no peribronchiolar inflammatory cuffs; alveolar spaces were uniform and open with thin septa; and arterioles showed no perivascular infiltrates, edema, or hemorrhage (Fig. 2F-H and Fig S4).

Infection with wild-type WU2 induced severe suppurative bronchopneumonia. Bronchioles displayed disruption of the pseudostratified columnar epithelium, dense neutrophilic cuffs (Fig. 2F, arrows), and occasional collapsed or shrunken lumens (Fig. S4). Alveolar regions showed extensive consolidation with dense localized neutrophilic infiltrates (Fig. 2G, arrows), fibrinous exudate (pink proteinaceous material), edema, inflammatory cell accumulation, and erythrocyte infiltration indicative of lung hemorrhage (Fig. 2G). Perivascular/arteriolar areas exhibited prominent neutrophilic cuffing and edema, reflecting strong cytotoxicity and vascular involvement (Fig. 2H). The WU2Δ*spxB* mutant exhibited severe but distinct pathology compared to wild-type WU2. In bronchioles, abundant inflammatory infiltrates (arrow) densely cuffed the airways, accompanied by swollen epithelial structures (asterisks) suggestive of goblet cell hyperplasia, mucin accumulation, or edematous changes, along with epithelial disruption and occasional collapsed/shrunken lumens (Fig. 2F). Alveolar regions showed persistent consolidation with inflammatory cell accumulation and variable edema, though less uniform than in wild-type (Fig. 2G and Fig. S4B). Perivascular/arterial areas displayed marked cuffing and inflammatory involvement around vessels (Fig 2H). These findings indicate robust local inflammation and airway remodeling in the absence of SpxB, potentially reflecting compensatory mechanisms or reduced acute cytotoxicity but sustained tissue damage.

Infection with WU2Δ*lctO* produced intermediate changes: bronchioles showed moderate epithelial disruption and neutrophilic cuffs; alveolar regions had consolidation and inflammatory cell gathering comparable to or slightly less than wild-type, with occasional hemorrhage; arteriolar/perivascular areas displayed intermediate cuffing and edema, aligning with the lesser contribution of LctO to total H₂O₂ production (Fig. 2F-H). The WU2Δ*spxB*Δ*lctO* double mutant induced pathology closely resembling wild-type WU2 across regions: bronchioles with prominent epithelial disruption, collapsed lumens, and dense cuffs; alveoli with severe neutrophilic consolidation, exudate, inflammatory cell accumulation, and erythrocyte extravasation representing hemorrhage; arterioles with marked perivascular neutrophilic infiltrates and edema (Fig. 2F-H). This restoration of severe suppurative and hemorrhagic features supports compensatory mechanisms that overcome the attenuation observed in the single Δ*spxB* mutant.

Collectively, these multi-regional histological findings demonstrate that the pneumococcal pyruvate node pathway—predominantly through SpxB, with a secondary contribution from LctO—drives acute cytotoxicity, bronchiolar epithelial disruption, neutrophilic inflammation, alveolar consolidation, perivascular involvement, and pulmonary hemorrhage at euthanasia in the ST3 WU2 model. The observed differences in pathology severity across the mutants underscore the critical role of pathway-specific metabolic regulation in determining the extent and distribution of tissue damage in the lower respiratory tract.

### SpxB-Dependent Reorganization of Bronchial Epithelial Membranes and Shedding of Bacterium-Laden Structures during Serotype 3 Pneumococcal Pneumonia

To examine the ultrastructural impact of SpxB-dependent pyruvate metabolism in the lower respiratory tract during ST3 pneumococcal pneumonia, we performed high-resolution confocal microscopy on lung tissues from mice intranasally infected with wild-type WU2, WU2Δ*lctO*, or WU2Δ*spxB*. Histological sections of the entire left lung were triple-labeled with DAPI (for nuclear DNA), WGA conjugated to Alexa Fluor 488 (WGA-AF488; staining GlcNAc/Neu5Ac-containing host plasma membranes and pneumococcal peptidoglycan), and an Alexa Fluor 555-conjugated anti-ST3 capsular polysaccharide antibody. Images were acquired and analyzed by high-resolution confocal microscopy and Imaris software, respectively.

Bronchioles from mock-infected mice exhibited intact pseudostratified columnar epithelium, with continuous apical and basolateral WGA staining and minimal background capsular signal (Fig. 3A, arrowheads). In contrast, wild-type WU2 infection induced prominent WGA-positive membrane protrusions extending from the apical surface of bronchial epithelial cells into the airway lumen (Fig. 3A, arrows). These protrusions frequently terminated in bulbous structures that co-localized with encapsulated pneumococci, indicating physical association of intact diplococci or short chains with host-derived membranes (Fig. 3A, dashed circles). Many bulbous structures appeared to be detaching, releasing 1–3 µm diameter vesicles containing ST3 capsule-positive pneumococci enveloped in host membrane (Fig. 3A, asterisks). Detached, bacterium-laden blebs were often observed deeper within bronchiolar tissue, suggesting a directed, metabolism-dependent mechanism that facilitates bacterial translocation.

**Fig. 3.**
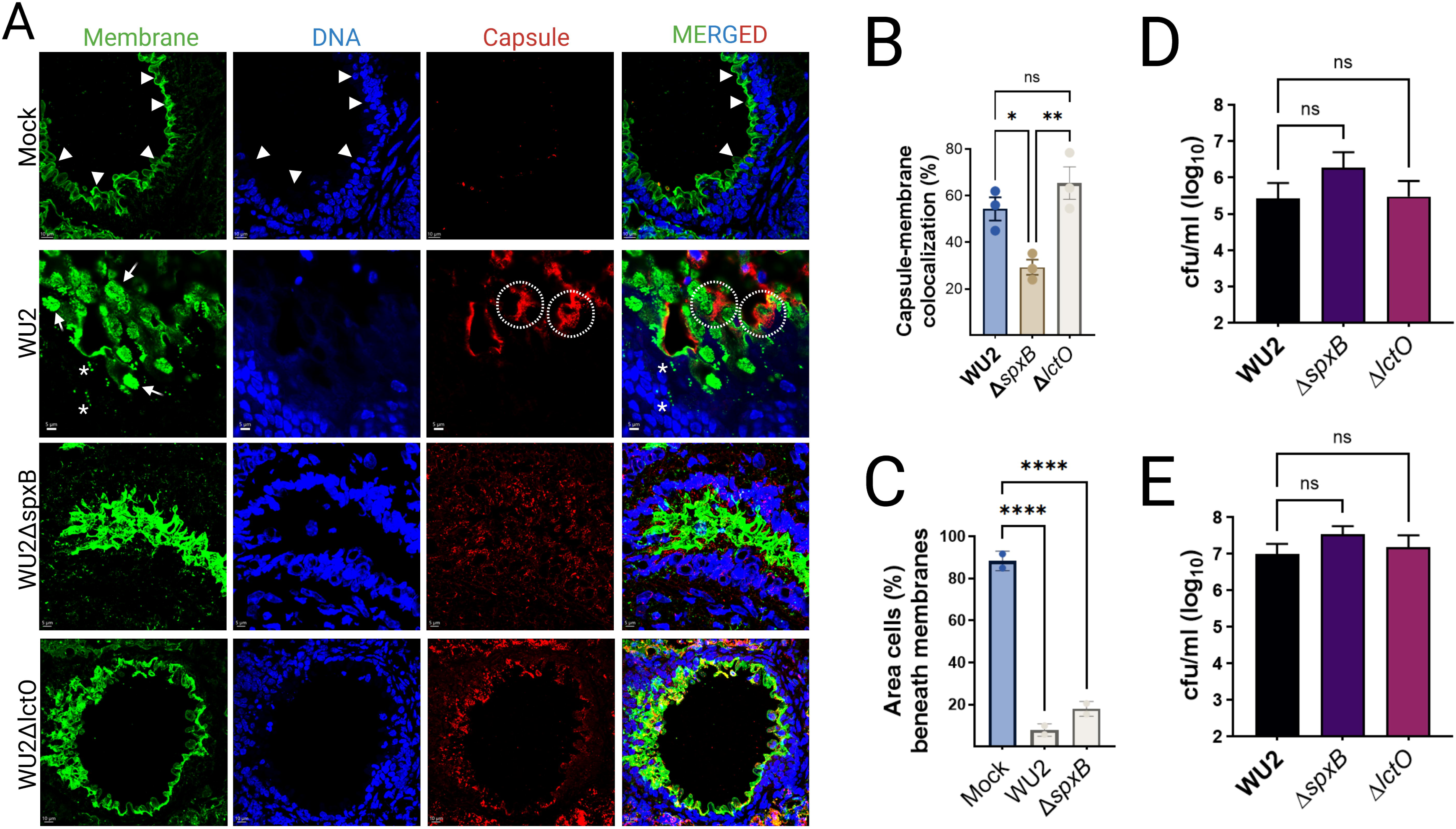
SpxB-dependent reorganization of bronchial epithelial membranes and shedding of bacterium-laden structures during serotype 3 pneumococcal pneumonia. (A) Representative confocal microscopy images of bronchiolar sections from mock-infected or infected C57BL/6 mice (endpoint). Lung tissues were triple-labeled with DAPI (blue; nuclear DNA), wheat germ agglutinin conjugated to Alexa Fluor 488 (WGA-AF488, green; staining GlcNAc/Neu5Ac residues on host plasma membranes), and Alexa Fluor 555-conjugated anti-serotype 3 capsular polysaccharide antibody (red; capsule). Mock-infected bronchioles show intact pseudostratified columnar epithelium (arrowheads). WU2-infected bronchioles exhibit prominent WGA-positive membrane protrusions/reorganization (arrows), bulbous structures (dashed circles), and vesicles (asterisks) co-localized with encapsulated pneumococci. Sections from WU2Δ*spxB*- or WU2Δ*lctO*-infected mice display marked architectural disruption and diffuse bacterial dissemination (lower panels). Scale bars: 10 µm (main images) or 5 µm (zoomed insets); representative of n=2–3 mice per group. (B) Quantitative analysis of capsule-membrane co-localization: percentage of capsular polysaccharide signal overlapping with WGA-positive membrane signal in infected bronchioles. Data are mean ± SE; statistical comparison by unpaired Student’s t-test (p < 0.05). (C) Quantitative analysis of epithelial membrane disruption: percentage of area beneath the bronchial epithelial cell layer occupied by detached/shed structures or spaces (expressed as % of total imaged area). Data are mean ± SE; statistical comparisons by one-way ANOVA with Dunnett’s multiple-comparisons test (***p < 0.0001 vs. mock). (D–E) In vitro adhesion assays of WU2 and mutant derivatives to human bronchial Calu-3 cells under (D) lung-like oxygen conditions (14% O₂) or (E) nasopharyngeal-like conditions (∼21% O₂). Bacterial adherence quantified as log₁₀ CFU/mL recovered after incubation and washing. Data are mean ± SE from triplicate wells; statistical comparisons by one-way ANOVA with Dunnett’s post-test (ns, not significant).

In bronchioles from mice infected with WU2Δ*spxB* or WU2Δ*lctO*, these organized apical membrane protrusions and bacterium-laden blebs were absent. Instead, the epithelium displayed marked architectural disruption and diffuse bacterial dissemination at euthanasia (Fig. 3A). The extensive membrane reorganization and luminal extrusion of bacterium-containing structures characteristic of wild-type WU2 infection were largely abolished in both mutants (Fig. 3A, lower panels), despite comparable bacterial burdens in the lungs at this time point (Fig. 2D). Quantitative analysis confirmed significantly greater co-localization of capsular polysaccharide and WGA-positive membrane signals in WU2-infected mice compared to those infected with WU2Δ*spxB* or WU2Δ*lctO* (Fig. 3B).

Epithelial cytotoxicity was assessed by quantifying the area (µm²) of nuclear pyknosis and fragmentation (i.e., absence of cell nuclei) immediately beneath the bronchial lining, as a proxy for cell detachment and cytotoxicity. Compared to mock-infected animals, nuclear condensation/fragmentation area within 5 µm of the WGA-defined epithelial border was significantly increased in all infected groups (****P < 0.0001 vs. mock), but levels were comparable across pneumococcal strains (Fig. 3C).

Given the striking differences in structural association with the bronchial lining between wild-type and pyruvate node mutants, we used polarized Calu-3 bronchial epithelial cells as a reductionist *in vitro* model to evaluate potential strain-specific changes in adhesion. No differences in adhesion to epithelial cells were observed among ST3 strains, regardless of growth under oxygen-limited (Fig. 3D) or oxygen-rich conditions (Fig. 3E).

Collectively, these findings demonstrate that functional pyruvate flux through SpxB (with supportive contribution from LctO) drives targeted reorganization of bronchial epithelial membranes. This process promotes the formation and extrusion of bacterium-laden blebs that encapsulate intact pneumococci, thereby facilitating organized bacterial penetration into underlying tissue in a manner independent of overt local cytotoxicity. These observations highlight a critical role for SpxB-mediated metabolic activity in orchestrating non-lytic host-pathogen interactions during ST3 pneumococcal pneumonia. In contrast, pyruvate node mutants exhibit dysregulated invasion, which correlates with the increased cytotoxicity observed in histopathological analyses.

### Disruption of alveolar and endothelial membrane integrity and enhanced inflammatory infiltration in oxidase-deficient serotype 3 pneumococcal pneumonia

To investigate the consequences of impaired pyruvate metabolism in distal lung compartments during ST3 pneumococcal pneumonia, alveolar parenchyma and pulmonary arteries were also examined using high-resolution confocal microscopy with the same triple-labeling protocol (DAPI for nuclear DNA, WGA-Alexa Fluor 488 for GlcNAc/Neu5Ac-containing host membranes and pneumococcal peptidoglycan, and Alexa Fluor 555-conjugated anti-ST3 capsular polysaccharide antibody).

In mock-infected mice, alveolar septa and endothelial linings exhibited continuous, uniform WGA staining, delineating intact structural barriers with minimal inflammatory cells and no capsular signal (Fig. 4A–B). Infection with wild-type WU2 resulted in moderate disruption of membrane continuity, evidenced by focal fragmentation and punctate WGA patterns in alveolar septa and vascular walls. Inflammatory infiltration was limited, primarily consisting of scattered cells in alveolar spaces, with ST3 capsular polysaccharide signal largely restricted to septal tissue and perivascular regions rather than freely distributed in the airspaces (Fig. 4A–B).

**Fig. 4.**
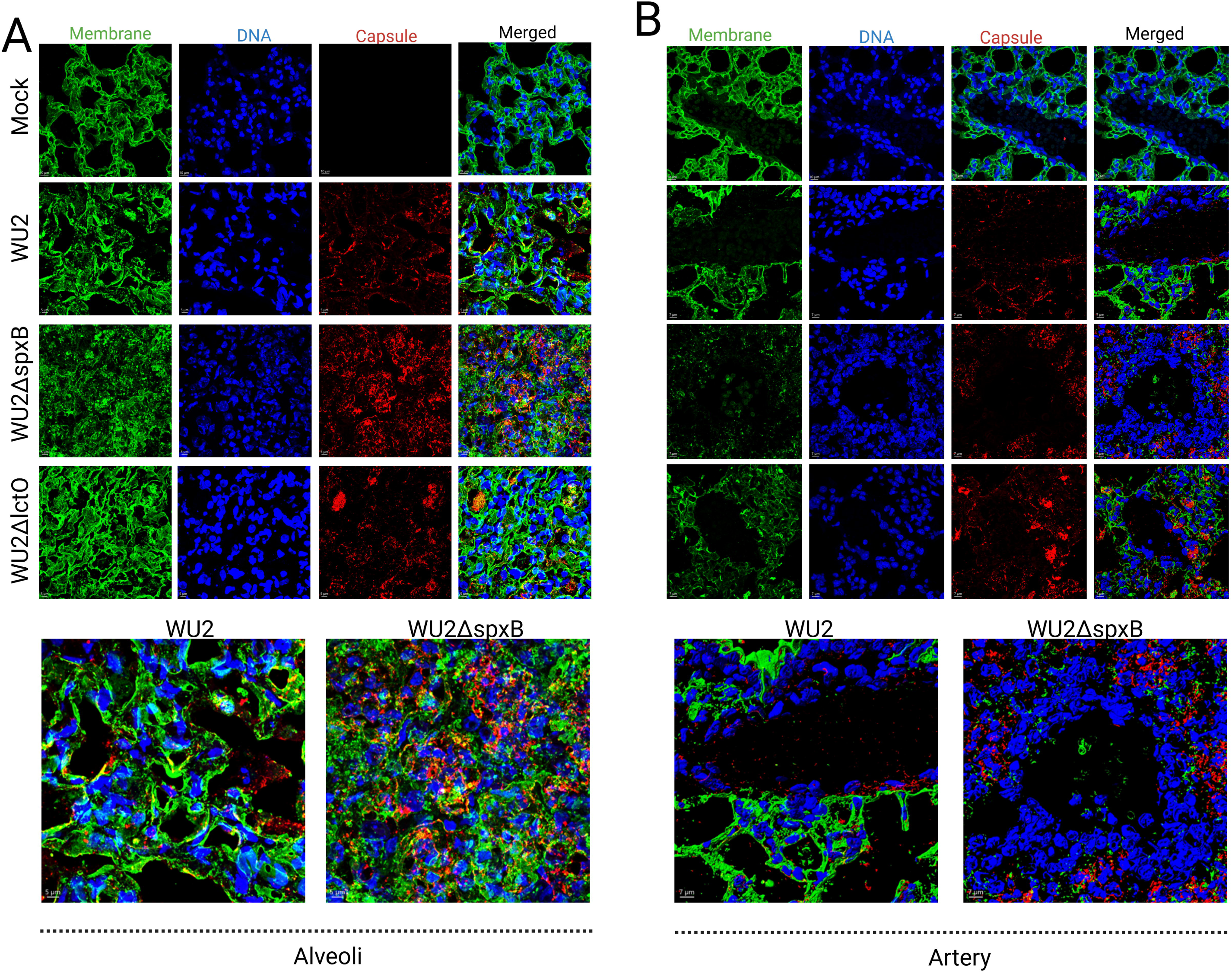
Disruption of alveolar and endothelial membrane integrity and enhanced inflammatory infiltration in oxidase-deficient serotype 3 pneumococcal pneumonia. (A) Representative confocal microscopy images of alveolar sections from mock-infected or infected C57BL/6 mice (experimental endpoint) with wild-type WU2, WU2Δ*spxB*, or WU2Δ*lctO*. Lung tissues were triple-labeled with DAPI (blue; nuclear DNA), wheat germ agglutinin conjugated to Alexa Fluor 488 (WGA-AF488, green; staining GlcNAc/Neu5Ac residues on host plasma membranes and pneumococcal peptidoglycan), and Alexa Fluor 555-conjugated anti-serotype 3 capsular polysaccharide antibody (red; capsule). Z-stack projections were acquired to visualize 3D architecture. Mock-infected alveoli display intact, thin alveolar septa with minimal WGA-positive membrane signal and no bacterial presence. WU2-infected alveoli show marked disruption: thickened septa, irregular membrane staining (green protrusions/edema), dense inflammatory infiltrates (blue DAPI+ cells), and encapsulated bacteria (red) associated with damaged structures. Sections from WU2Δ*spxB*- and WU2Δ*lctO*-infected mice exhibit similar but variably attenuated pathology, with persistent membrane disruption and bacterial dissemination in some fields. Bottom panels show zoomed micrographs of representative regions for each strain to highlight details of alveolar damage and bacterial-host interactions. (B) Representative confocal microscopy images of pulmonary artery/arteriole sections, processed and labeled identically to (A). Mock-infected vessels show smooth endothelial lining (green WGA) with intact architecture. WU2-infected vessels display endothelial membrane disorganization, perivascular edema/inflammation (dense DAPI+ infiltrates), and encapsulated pneumococci adherent to or within disrupted endothelium. Mutant-infected sections (Δ*spxB* and Δ*lctO*) show comparable endothelial disruption and inflammatory cuffing, with bacteria present in vascular/perivascular spaces. Bottom panels are zoomed views of corresponding strains to emphasize vascular pathology. Scale bars are indicated in the bottom-left corner of each panel (e.g., 5–7 µm for zoomed insets; larger for overview fields). Images are representative of n=3–5 mice per group, with z-stacks used for depth-resolved analysis.

Infection with WU2Δ*spxB* displayed severe and widespread membrane damage, characterized by extensive loss of continuity, highly fragmented and punctate WGA staining throughout the alveolar parenchyma and endothelial layers. This was accompanied by markedly increased inflammatory cell accumulation (dense clusters in alveolar spaces and interstitium) and intense, diffuse capsular polysaccharide staining filling the alveolar lumen, interstitium, and vascular areas (Fig. 4A–B). In contrast, WU2Δ*lctO*-infected lungs showed intermediate membrane disruption, with fragmentation and punctate WGA patterns more pronounced than in wild-type but less severe than in WU2Δ*spxB*. Capsular signals appeared aggregated and enclosed in localized clusters rather than diffusely disseminated, and inflammatory infiltration was comparable to wild-type levels, with fewer dense cellular clusters relative to WU2Δ*spxB* (Fig. 4A–B).

These patterns occurred despite equivalent overall bacterial burdens in the lungs across groups at this time point (Fig. 2D). The findings indicate that ablation of SpxB-dependent pyruvate metabolism leads to the most severe alveolar and endothelial barrier disruption, uncontrolled bacterial dissemination, and amplified inflammatory responses, whereas LctO disruption results in a milder, intermediate phenotype with aggregated bacterial localization and restrained infiltration. Collectively, functional pyruvate flux through SpxB and LctO modulates the extent of structural damage, bacterial spread, and host inflammatory dynamics in distal lung compartments.

### RNA Sequencing reveals immune dysregulation in pyruvate oxidase-deficient pneumococcal lung infection

To extend histopathological and high-resolution confocal microscopy findings indicating dysregulated immune responses in the absence of key pyruvate metabolic pathways, we performed bulk RNA sequencing on lung tissues from mice infected with ST3-Spn strains (wild-type WU2, WU2Δ*spxB*, or WU2Δ*lctO*) or mock-infected controls. Lungs were harvested at 48 h post-infection. A total of 35,566 genes were annotated, of which 16,105 were expressed and included in differential expression analyses. Comparisons revealed 2,783 DEGs between mock- and wild-type WU2-infected lungs, 2,890 between mock- and WU2Δ*spxB*-infected lungs, and 338 between wild-type WU2- and WU2Δ*spxB*-infected lungs. Of these, 1,689, 1,860, and 125 genes were significantly upregulated, while 1,094, 1,030, and 213 were significantly downregulated, respectively (FDR < 0.05, log_2_ fold-change >1).

Functional enrichment analysis using g:Profiler against the Gene Ontology (GO) database revealed significant enrichment of genes associated with oxidoreductase activity (GO:0016705). In WU2Δ*spxB*-infected lungs versus wild-type WU2 infected lungs, multiple genes in this category were significantly downregulated (log_2_ fold-change ≥ 1; FDR < 0.05), consistent with reduced host oxidative stress due to the absence of SpxB-mediated metabolic pathway (Fig. 5A). Notably, *nos2* (encoding inducible nitric oxide synthase, iNOS) was significantly downregulated in WU2Δ*spxB*-versus wild-type WU2-infected lungs. Several cytochrome P450-related genes were downregulated in wild-type WU2-infected lungs versus mock-infected controls, consistent with epithelial injury and/or metabolic suppression (Fig. 5A). This dysregulation of cytochrome P450-related genes was attenuated in the absence of SpxB. For example, *cyp2e1* (encoding a cytochrome P450 family monooxygenase) showed strong downregulation in wild-type WU2-infected lungs versus WU2Δ*spxB* (log₂ fold-change = –4.75), moderate downregulation in wild-type WU2-infected lungs versus mock (log₂ fold-change = –2.47), and upregulation in WU2Δ*spxB*-infected lungs versus mock (log₂ fold-change = +2.28) (FDR < 0.05 for all).

**Fig. 5.**
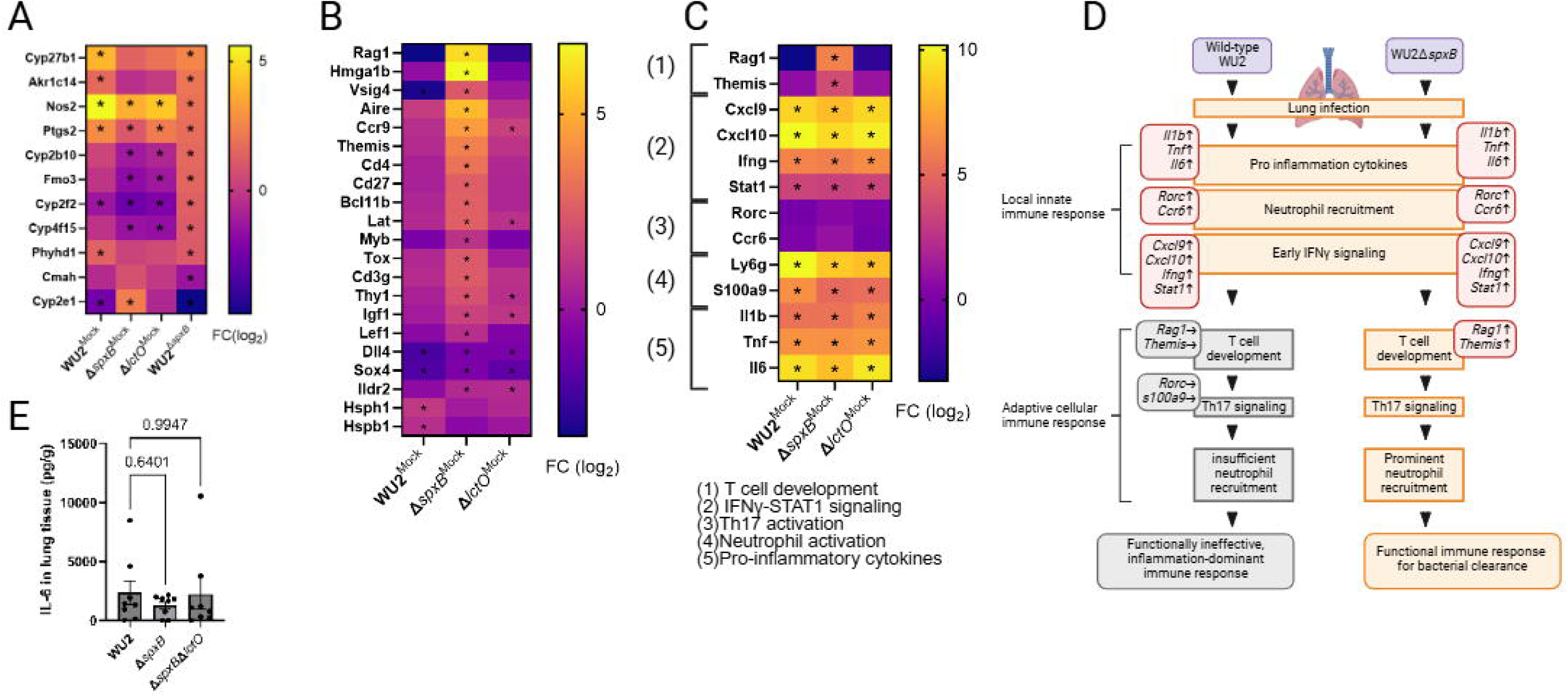
Immune dysregulation in pyruvate oxidase-deficient pneumococcal lung infection in RNA Sequencing. (A, B, C) Bulk RNA-seq analysis of the lung tissues from mock-infected mice or mice infected for 48 h with the indicated strain. Heatmaps depict relative expressions (log2 fold change) of selected genes associated with (A) oxidoreductase-related processes, (B) selected biological process of T cell activation, and (C) additional immune-related processes with group information. (D) Graphical summary of the difference in host immunological response to wild-type WU2 and WU2Δ*spxB*. The upward and downward arrows represent upregulation and downregulation of the gene, respectively. (E) IL-6 quantification in the mouse lung tissue by enzyme linked immunosorbent assay. Data in the panel is shown as mean ± SE. Statistical analyses were performed using one-way ANOVA with Dunnett’s multiple-comparisons test.

Notably, 15 genes encoding for proteins directly involved in the biological process of T cell activation (GO:0042110) including *Rag1*, and *Themis*, were upregulated in lung tissue from mice infected with WU2Δ*spxB* compared with the mock-infected tissue (log₂ fold-change 1.0 – 6.8; FDR < 0.05), but not in the tissue infected with wild-type (log₂ fold-change -3.2 – 1.4; FDR ≥0.05) (Fig. 5B). In addition, several genes, such as *Vsig4, Dll4*, and *Sox4* related to acute inflammatory response, were downregulated in wild-type WU2-infected tissues compared with the mock-infected tissue (log₂ fold-change -1.8 – -3.0; FDR < 0.05). Furthermore, the genes such as *Rorc* and *Ccr6* involved in Th17 activation were not dysregulated (log₂ fold-change -0.2 – 0.1; FDR ≥0.05). In both wild-type and mutant infections, genes linked to innate immune response such as pro inflammatory cytokines (e.g., *il6*, *tnf*), neutrophil activation, and early phase of IFNγ-STAT1 signaling were highly upregulated (Fig. 5C, D). Consistent with these findings, ELISA measurements revealed comparable elevated IL-6 levels in lung tissue from mice infected with wild-type WU2 or pyruvate node mutants (Fig. 5E). These results suggest that SpxB-dependent pyruvate node activity in wild-type WU2 suppresses T cell activation and acute inflammatory pathways in the infected lung (Fig. 5D). This gene expression signature aligns with the histological evidence of bacterium-laden blebs along the bronchiolar epithelium in WU2-infected mice, potentially representing a pathway-mediated mechanism that sequesters pneumococci and limits their exposure to host immune surveillance.

## Discussion

In the present study, we demonstrate that the ST3 strain WU2 exploits the SpxB-dependent pyruvate metabolic pathway to facilitate invasion of the bronchiolar epithelium. This process involves the formation of pneumococci-laden membrane blebs that incorporate host cell membrane components, enabling both epithelial invasion and sequestration of the bacteria from host immune surveillance. Consequently, this mechanism appears to attenuate the adaptive immune response, as evidenced by our transcriptomic analysis showing suppression of T cell activation and acute inflammatory pathways in wild-type WU2-infected lung tissue compared with the WU2Δ*spxB* mutant. Strong corroborative evidence for this model derives from *in vitro* oxygen-sensitivity assays, which revealed a pronounced fitness cost in ST3 strains, including wild-type WU2, under lung-relevant oxygen concentrations (∼21% O₂, PO₂ ∼160 mmHg in ambient/upper airway air). In contrast, exposure to lower lung-relevant oxygen levels (∼14% O₂ equivalent, PO₂ ∼100–105 mmHg in alveoli) triggered a distinct metabolic shift compared with growth under pharyngeal conditions, consistent with oxygen-dependent regulation of the pyruvate node and SpxB activity. SpxB, the pneumococcal pyruvate oxidase, catalyzes the aerobic conversion of pyruvate to acetyl phosphate, CO₂, and H₂O₂, providing ATP while generating reactive oxygen species that can influence both bacterial fitness and host interactions.

The immune response to lung infection is initiated by innate recognition of invading bacteria, primarily by alveolar macrophages, which sense Spn and induce proinflammatory cytokines such as TNF-β, IL-1β, and IL-6 to drive local immune responses and recruit neutrophils [35, 36]. Subsequent adaptive T cell immunity, including Th17 and Th1, sustains neutrophil and macrophage activation and contributes to bacterial clearance [37]. However, ST3 strains such as WU2 appear to subvert this cascade. A previous *in vivo* study demonstrated that ST3-infected mice exhibited a more exuberant inflammatory response in the lung, higher levels of IFN-γ-inducible gene expression compared to ST8-infected mice, and reduced ST3 virulence in CXCR3-deficient mice, suggesting a role for IFN-γ-inducible chemokines (e.g., CXCL9/10/11) driven recruitment and pathogenesis [24]. Furthermore, ST3-Spn evades IL-17 mediated neutrophil killing, leading to persistent neutrophil accumulation in the lung and consequent neutrophil-mediated lung damage; this pathology is attenuated upon neutrophil depletion [25]. Our findings extend this paradigm by linking SpxB activity to immune attenuation: the oxygen-triggered metabolic shift in the lung environment promotes bleb-mediated invasion and sequestration while dampening T cell priming and acute inflammation. This SpxB-dependent, responsive to the physiological oxygen gradient from the upper (∼21% O₂, favoring colonization) to lower respiratory tract, may allow ST3 to orchestrate a delicate balance: triggering robust initial innate hyper-responsiveness (e.g., neutrophil influx and cytokine storm) while suppressing effective adaptive immune clearance (e.g., via T cell activation dampening and sequestration in blebs). This dual modulation likely underpins the known hypervirulence of ST3 pneumococci and their propensity for prolonged persistence and tissue damage in the lower respiratory tract.

In support of the hypotheses above, our transcriptome studies revealed that infection with wild-type WU2 and pyruvate node mutants induced significant upregulation of genes encoding proinflammatory cytokines (e.g., *Il1b*, *Il6*, and *Tnf*) and mediators of neutrophil recruitment (e.g., *ly6g*, and *S100a9*) [38, 39]. This pattern correlated with the similar levels of IL-6 protein measured by ELISA in lung homogenates of infected mice. Similarly, all strains triggered robust transcription activation of IFN-γ signaling pathway genes, such as *Cxcl9*, *Cxcl10* and *Ifng* [40]. This coordinated activation of neutrophils- and macrophages-related genes strongly correlates with the severe histopathological findings, including the dense inflammatory infiltrate in bronchiole, alveoli and pulmonary arteries as well as pronounced ultrastructural changes (e.g., alveolar damage, vascular involvement, and tissue remodeling) identified by high resolution confocal microscopy.

In agreement with the limited IL-17–mediated bacterial clearance observed in ST3 infections, mice infected with any WU2 strain did not induce transcriptional activation of genes *Rorc* and *Ccr6*, whose products are central to IL-17 pathway activity and Th17 differentiation [41, 42].

Remarkably, disrupting the SpxB-mediated pyruvate node pathway in strain WU2 led to enhanced activation of T cell responses in the lung, as evidenced by strong differential upregulation of genes implicated in T cell activation and maturation, including *Rag1* and *Themis*. These findings suggest that ST3 strains, such as WU2, actively limit adaptive T cell immunity during pulmonary infection (Fig. 5D), potentially through SpxB-dependent mechanisms such as metabolic shifts or bleb-mediated sequestration (i.e., bulbous structures in WU2-infected bronchioles) that dampen effective antigen presentation or T cell priming [43, 44]. In support of this model, upregulation of the canonical receptor *Cxcr3* was not observed (data not shown), which is consistent with exaggerated innate inflammation in the absence of effective T cell recruitment, likely due to evasion of ST3 pneumococci from antigen-presenting cells.

Pneumococcus exhibits distinct behaviors in response to environmental changes, but these responses vary among strains and serotypes [45]. Oxygen levels can modulate capsule production, virulence-related gene expression, and metabolic reprograming[46–48]. A previous dual RNA-seq study in a murine pneumonia model infected with ST3 revealed upregulation of bacterial virulence genes in the in pleural fluid, a relative hypoxic environment[26]. Our growth kinetic assays demonstrated a unique sensitivity to H₂O₂-mediated stress in ST3 strains, including WU2, suggesting that the relatively lung hypoxic environment provides a favorable niche for this serotype. This adaptation likely enables metabolic reprograming of growth and virulence gene transcription. Such oxygen-dependent fitness advantages may help explain the strong clinical association of ST3 with severe infections in low-oxygen niches, such as empyema and acute otitis media [7, 49].

Furthermore, this unique oxygen-sensitive growth disadvantage in ST3 was mitigated by *spxB* mutation, which clearly showed that SpxB-dependent pyruvate flux impairs WU2 fitness in high-oxygen environments (e.g., nasopharyngeal/pharyngeal conditions, ∼21% O₂). Alternations in bacterial metabolic pathways are known to influence biological behaviors[50, 51]. The observed fitness rescue in the mutant may involve compensatory metabolic changes beyond direct SpxB-derived H₂O₂-mediated stress [52, 53].

We also observed distinct histopathological outcomes in lung tissues infected with wild-type WU2 and the WU2Δ*spxB* mutant, despite comparable bacterial colonization densities. In contrast to previous reports with other serotypes [16, 54], SpxB-deficient mutant disseminated diffusely within lung compartments and caused unstructured epithelial damage, highlighting the potential cytotoxicity of ST3 pneumococci as previously demonstrated [24]. Although wild-type WU2 also induced severe tissue damage, it promoted bronchiolar membrane remodeling, aggregated within membrane-bulbous structures, and localized preferentially in septal tissue and perivascular regions rather than freely distributing in the alveolar airspace, features not observed in our prior studies infecting with strain TIGR4 (serotype 4) or D39 (serotype 2) [16]. These unique behaviors are consistent with escape from the oxidative stress and host immune pressure, as supported by our growth kinetics and the gene expression analyses, and enables ST3-Spn to remain within tissues like intracellular bacteria [55], where they may be targeted by Th1-IFNγ-macrophage axis while evading adaptive cellular immune system, ultimately leading to severe tissue disruption[24, 25].

An additional important finding was that the bacterial burden of WU2Δ*spxB* in the upper respiratory tract was markedly lower than that of wild-type WU2 in the mouse pneumonia model. Previous murine studies using Δ*spxB* mutants of other serotypes also reported reduced bacterial burdens in nasopharynx compared to the wild-type strains[56–58]. However, those differences were generally less pronounced, with mutant strains often maintaining substantial colonization levels. In contrast, the absence of SpxB activity in the ST3 background resulted in near-sterilizing defects in colonization. Although the precise mechanisms remain to be elucidated, this more severe phenotype in ST3 may reflect a heightened dependence on SpxB for evading host T-cell-mediated immune responses, suggesting a potentially serotype-specific role for this pathway. Further mechanistic studies are warranted and are currently ongoing in our laboratory.

In summary, ST3 pneumococci exploit the SpxB-driven pyruvate node pathway to induce host membrane-bleb formation in bronchiolar epithelial cells and limit adaptive T cell immunity, thereby facilitating invasion and persistence. A unique sensitivity to H₂O₂-mediated stress further supports metabolic adaptation to the relatively hypoxic lung environment. These SpxB-related, oxygen-dependent bacterium-host interactions represent a novel hallmark of ST3 pathogenesis, providing new insights and potential therapeutic targets.

## Acknowledgements

This study was supported in part by grants from the NIH, including R01AI175461 (to J.E.V.). Confocal microscopy studies were supported by the National Institute of General Medical Sciences (NIGMS) under Award Number P20GM121334. J.E.V. is also supported by NIGMS through the Molecular Center of Health and Disease (P20GM144041). KT and some experiments in this study are supported by the Robert Austrian Research Award by the International Society of Pneumonia and Pneumococcal Diseases and Pfizer. RNA-seq data analysis was performed at the UMMC Molecular and Genomics Facility, supported by funds from NIGMS (P20GM144041), Mississippi INBRE (P20GM103476), the Obesity, Cardiorenal, and Metabolic Diseases COBRE (P30GM149404), and the Mississippi Center of Perinatal Research (P20GM121334). The content is solely the responsibility of the authors and does not necessarily represent the official views of the NIH. The authors thank Dr. David Brown (UMMC) for his assistance with confocal microscopy.

## Author contributions

K.T. - Formal analysis, Investigation, Methodology, Project administration, Writing - original draft, Writing - review & editing. A.G.J.V. - Investigation, Supervision, Writing - review & editing. J.E.V. - Conceptualization, Data curation, Formal analysis, Funding acquisition, Investigation, Supervision, Writing - review & editing.

---

**Fig. S1. Growth kinetics of Streptococcus pneumoniae types under varying oxygen conditions.** (A) Growth parameters of serotype 3 strain WU2 compared to TIGR4, D39, and EF3030 under nasopharyngeal conditions (21% O₂) over 24 h. Bars represent mean values of lag time (h), maximum growth rate (Max V, 1/h), time to maximum growth rate (t at Max V, h), and maximum optical density (Max OD₆₀₀). (B) Comparative analysis of WU2 growth parameters relative to TIGR4, D39, and EF3030 under lung conditions (14% O₂) versus nasopharyngeal conditions, showing enhanced growth kinetics. (C) WU2 growth parameters under lung conditions (14% O₂) compared to TIGR4, D39, and EF3030. Data in all panels are shown as mean ± SE. Statistical analyses in panels were performed using one-way ANOVA with Šidák’s multiple comparisons test.

**Figure S2. Hydrogen peroxide production by Streptococcus pneumoniae strain WU2 and mutant derivatives under nasopharyngeal- and lung-mimetic oxygen conditions.** Wild-type WU2 and its isogenic mutant derivatives (Δ*spxB*, Δ*lctO*, Δ*spxB*Δ*lctO*, and Δ*spxB*Ω*spxB*) were inoculated in Todd-Hewitt broth supplemented with 0.5% yeast extract and incubated for 6 h at 37°C under either nasopharyngeal-mimetic conditions (21% O₂ + 5% CO₂) or lung-mimetic conditions (14% O₂ + 5% CO₂). Culture supernatants were harvested, filter-sterilized, and extracellular hydrogen peroxide (H₂O₂) concentration was quantified using the Amplex Red® hydrogen peroxide/peroxidase assay kit. Data are presented as mean ± SE from three independent biological replicates. Statistical comparisons were performed using one-way ANOVA followed by Šidák’s multiple-comparisons test. ns, not significant; *p < 0.05; **p < 0.01; ***p < 0.001; ****p < 0.0001 (exact significance levels are indicated in the figure panels where applicable).

**Figure S3. Growth kinetics of WU2, isogenic mutants, and *spxB*-complemented strain under varying oxygen conditions, and effects of catalase supplementation**. Wild-type WU2, WU2Δ*spxB*, WU2Δ*lctO*, WU2Δ*spxB*Δ*lctO*, and the *spxB*-complemented strain (WU2Δ*spxB*Ω*spxB*) were inoculated in Todd-Hewitt broth supplemented with 0.5% yeast extract (THY) and incubated at 37°C under controlled oxygen conditions. Optical density at 600 nm (OD₆₀₀) was measured every 20 min over 24 h. (A) Growth curves (OD₆₀₀ vs. time) of ST3 strains under nasopharyngeal-mimetic conditions (21% O₂). (B–E) Growth parameters derived from the curves in (A): (B) lag time (h), (C) maximum growth rate (Max V, h⁻¹), (D) time to maximum growth rate (t at Max V, h), and (E) maximum optical density (Max OD₆₀₀). (F–I) Growth parameters of the ST3 strains under lung-mimetic conditions (14% O₂): (F) lag time (h), (G) maximum growth rate (Max V, h⁻¹), (H) time to maximum growth rate (t at Max V, h), and (I) maximum optical density (Max OD₆₀₀). (J) Growth curves (OD₆₀₀ vs. time) for wild-type WU2, WU2Δ*spxB*, and WU2Δ*spxB*Δ*lctO* cultured under nasopharyngeal conditions (21% O₂) in THY, with or without addition of 100 U/mL catalase. Data in all panels represent mean ± SE from at least three independent biological replicates. Statistical comparisons were performed using one-way ANOVA followed by Šidák’s multiple-comparisons test. ns, not significant; exact significance levels are indicated in the figure panels where applicable.

**Fig. S4. Histopathological findings in the lungs of mice infected with serotype 3 *Streptococcus pneumoniae*.** (A–C) Representative hematoxylin and eosin-stained lung sections of the left lungs from mock-infected (PBS) or mice infected with Wild-type WU2, WU2Δ*spxB*, WU2Δ*lctO*, or WU2Δ*spxB*Δ*lctO* at the experimental endpoint. (A) Bronchioles, (B) Alveoli, and (C) Pulmonary arteries/arterioles. Images are representative of n=3–5 mice per group.

