## Supplementary figures and images for "Metabolic Rewiring at the Pyruvate Node Drives Severe Pneumonia and T-Cell Suppression in Serotype 3 *Streptococcus pneumoniae* Infection"

### Supplemental Figure S1

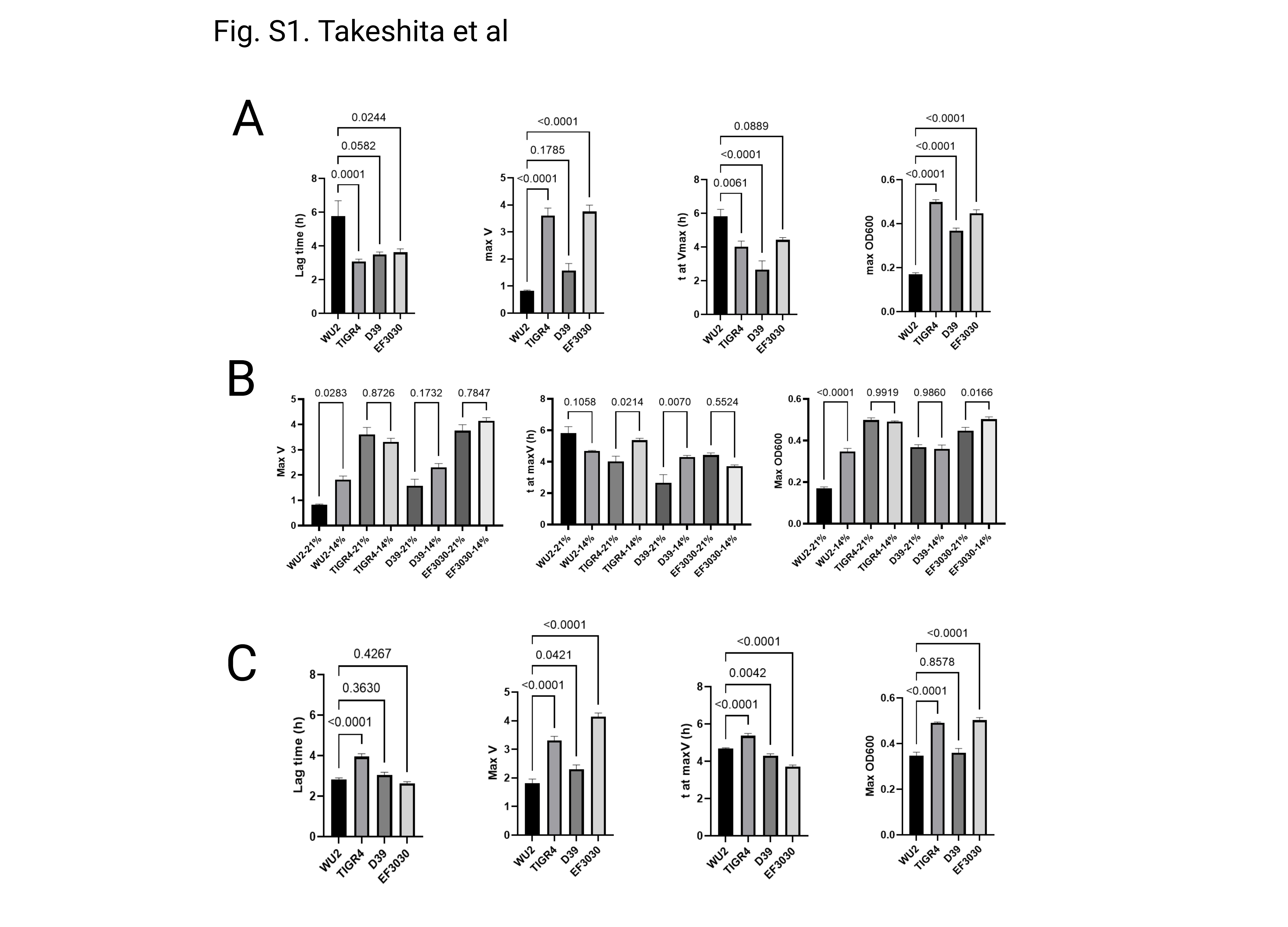

### Supplemental Figure S2

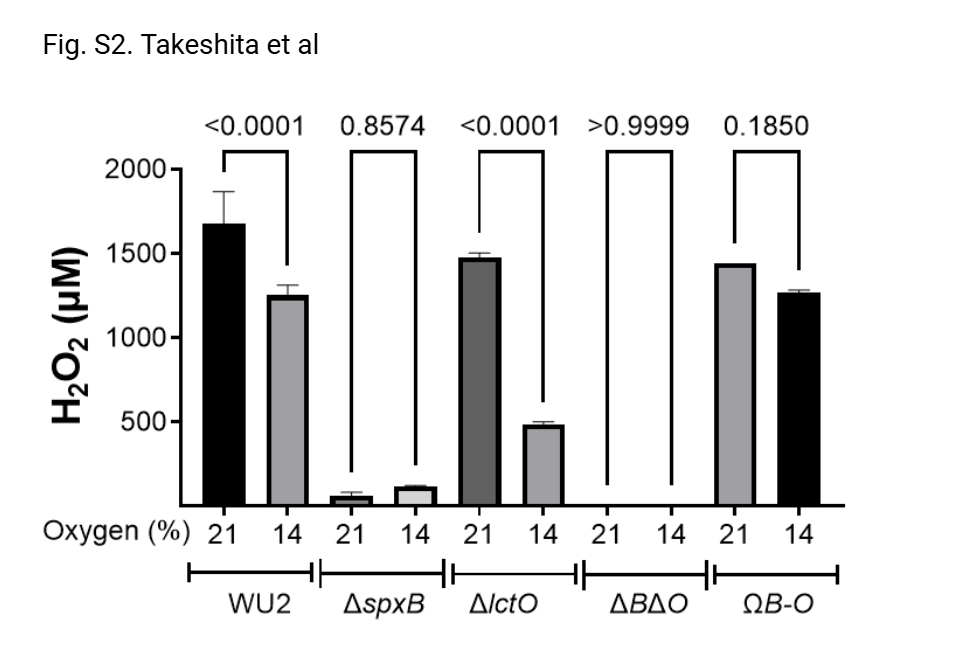

### Supplemental Figure S3

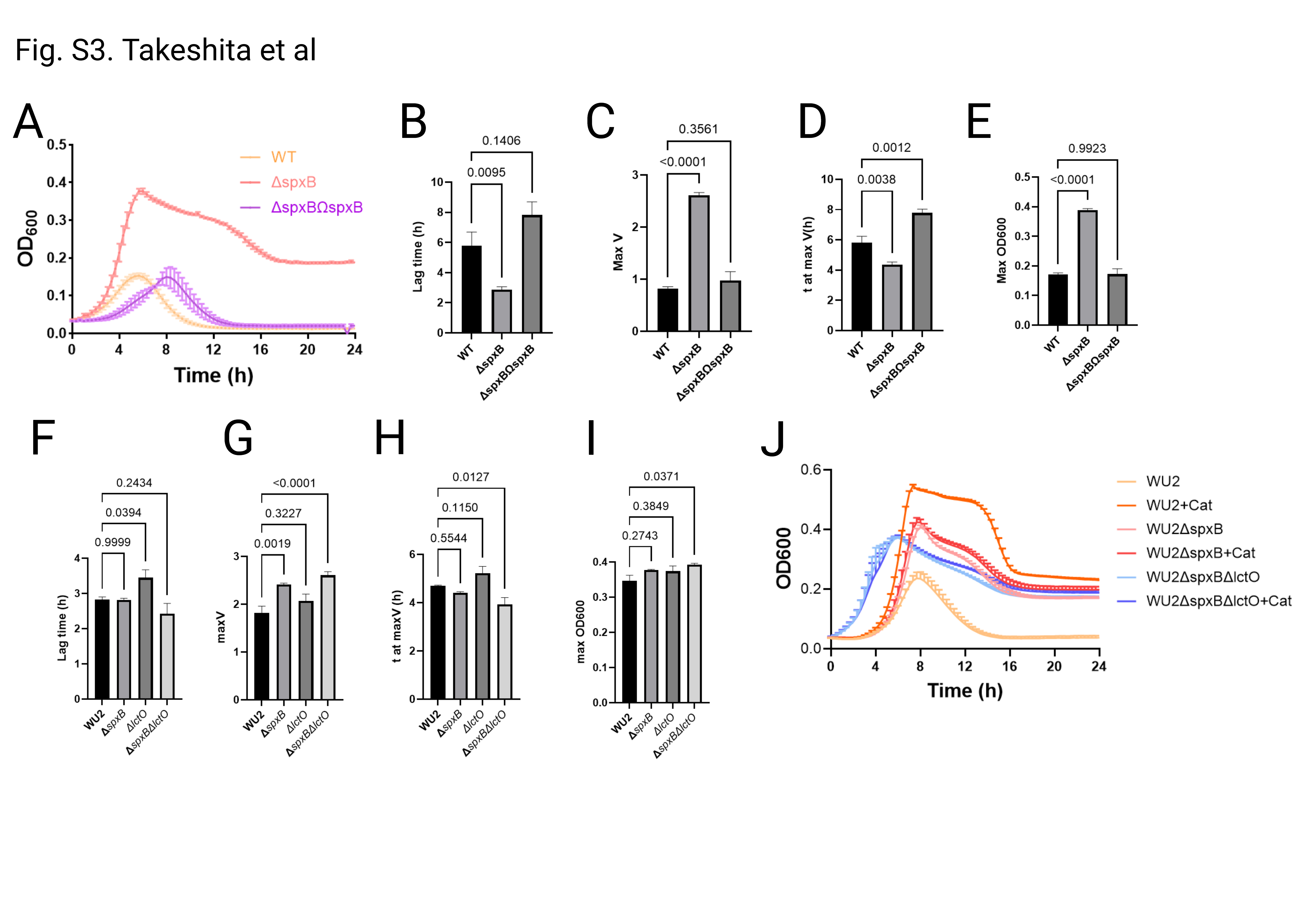

### Supplemental Figure S4

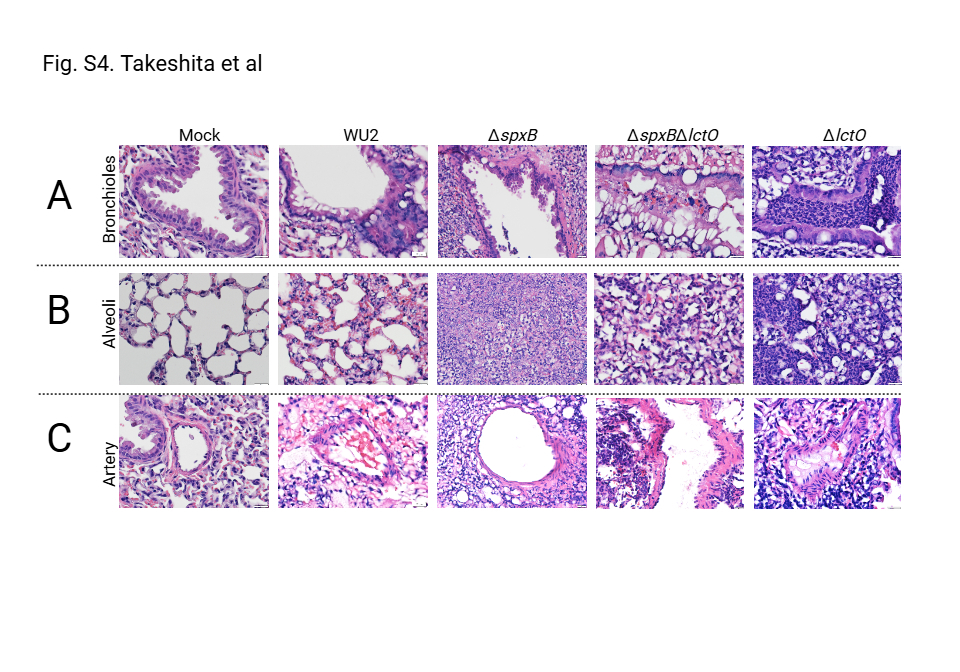
